# Structural basis for host recognition and superinfection exclusion by bacteriophage T5

**DOI:** 10.1101/2022.06.28.497910

**Authors:** Bert van den Berg, Augustinas Silale, Arnaud Baslé, Sophie L. Mader, Syma Khalid

## Abstract

A key but poorly understood stage of the bacteriophage life cycle is the binding of phage receptor binding proteins (RBPs) to receptors on the host cell surface, leading to injection of the phage genome and, for lytic phages, host cell lysis. To prevent a secondary viral infection by the same or a closely related phage, superinfection exclusion (SE) proteins can prevent the binding of RBPs via modulation of the host receptor structure in ways that are also unclear. Here we present the cryo-EM structure of the phage T5 outer membrane (OM) receptor FhuA in complex with the T5 RBP pb5, and the crystal structure of FhuA complexed to the OM SE lipoprotein Llp. Pb5 inserts four loops deeply into the extracellular lumen of FhuA and contacts the plug, but does not cause any conformational changes in the receptor, supporting the view that DNA translocation does not occur through the lumen of OM channels. The FhuA-Llp structure reveals that Llp is periplasmic and binds to a non-native conformation of the plug of FhuA, causing the inward folding of two extracellular loops via “reverse” allostery. The inward-folded loops of FhuA overlap with the pb5 binding site, explaining how Llp binding to FhuA abolishes further infection of *E. coli* by phage T5, and suggesting a mechanism for SE via the jamming of TonB-dependent transporters by small phage lipoproteins.

## Introduction

The increasing threat posed by multidrug-resistant bacteria, coupled to the lack of novel antibiotics, has led to a resurgent interest in the potential use of phage therapy to treat bacterial infections^1,2^, including phage steering^3^. Notwithstanding the enormous variety in phage structure and function, a defining moment during the infectious cycle of any phage is the high-affinity binding to protein and/or non-protein receptors on the host cell surface by phage receptor binding proteins (RBPs)^4^. This causes absorption of the phage on the cell surface and leads to the injection of the phage genome into the bacterial cell via a sequence of events that is still poorly understood. For lytic phages, genome injection leads to the assembly of progenitor phage in the host cytoplasm and cell lysis. To prevent the non-productive absorption of phage particles to already infected cells and post-lysis cell fragments, many phages express superinfection exclusion (SE) proteins early during infection^5^. One way to achieve SE is by inactivating the target receptor for RBPs. Similar to host protein receptor binding by RBPs, the mechanism by which SE proteins inactivate those receptors has not yet been visualised for any phage.

The lytic bacteriophage T5 is one of the model T coliphages that have been studied in great detail and which have stood at the basis of many fundamental discoveries in molecular biology^6-8^. Phage T5 is a caudal (tailed) virus within the family Demerecviridae and was sequenced in 2005 (ref. 9). Of its 162 predicted open reading frames^9^, more than half lack similarity to known genes and many T5 proteins are still uncharacterised, a common theme in phage biology. The overall morphology of T5 has been visualised via cryogenic electron microscopy^10^. As a Siphovirus, T5 has a long and flexible, non-contractile tail of ∼160 nm long, composed of the tail tube protein pb6 (ref. 11) and containing the tape measure protein pb2 that most likely perforates the *E. coli* cell envelope (Fig. 1a)^12^. The distal tail tip connects to a baseplate that anchors the three lateral tail fibers (LTF)^13^ composed of pb1 that bind LPS O-antigen reversibly^14^. The baseplate is composed of the Distal tail tip protein pb9 and the baseplate hub protein (BHP) pb3 that leads to the central straight fiber pb4 (Fig. 1a). Finally, the RBP pb5 (*oad* gene; Uniprot P23907) mediates irreversible T5 absorption to *E. coli* cells^15^ and is likely located at the distal end of pb4 (Fig. 1a). With exception of the monomeric RBP pb5, all tail proteins likely form oligomers within the intact phage^13^.

**Figure 1.**
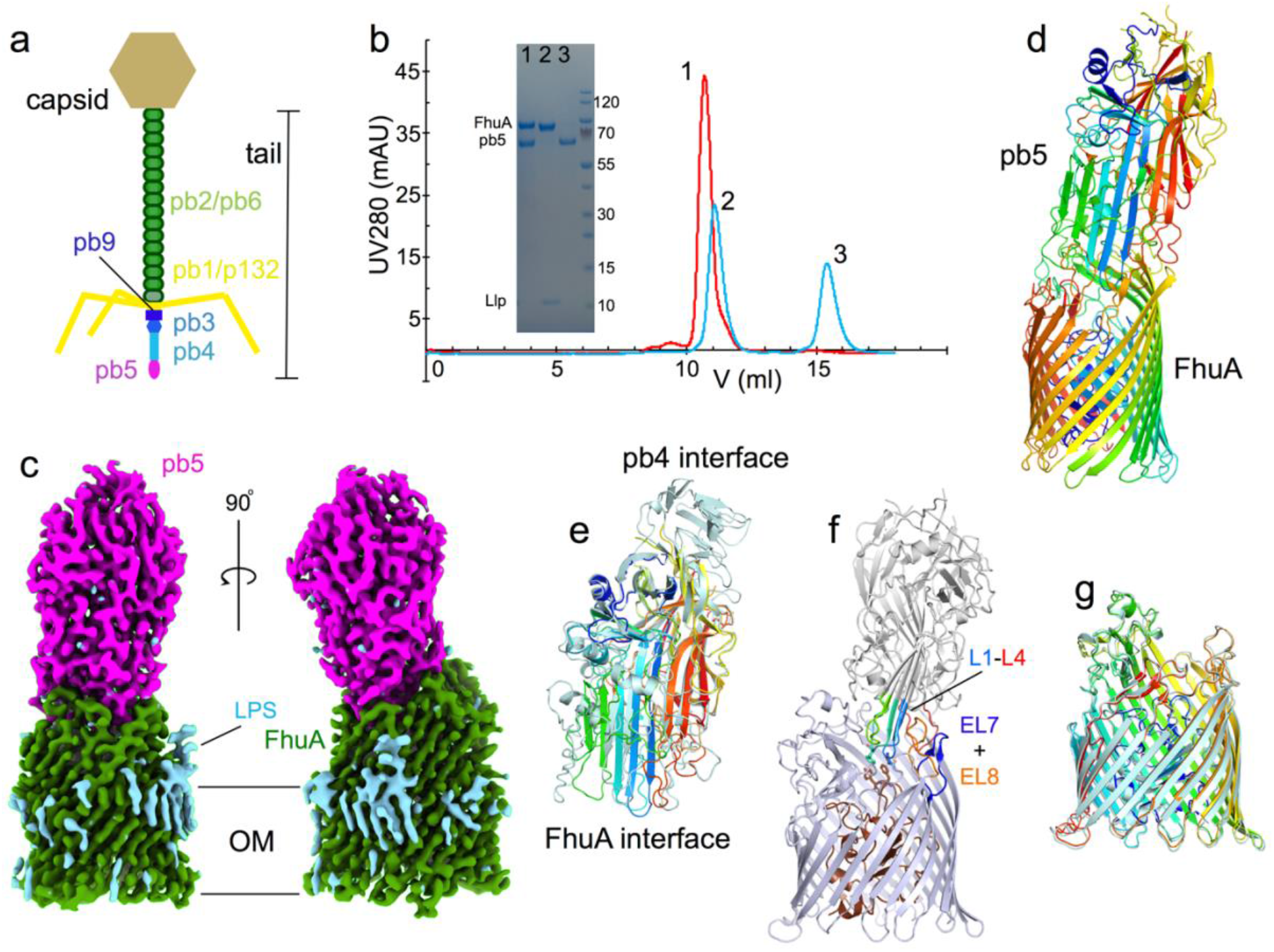
Cryo-EM structure of the FhuA-pb5 complex. **a**, Schematic representation of bacteriophage T5. **b**, Analytical SEC profiles for samples containing FhuA + pb5 (red) and FhuA-Llp + pb5 (cyan). One nmole of each protein was used. Peaks are numbered and were analysed via SDS-PAGE (inset). Band identities are shown on the left and the size of molecular weight markers on the right. Curves shown are representative for three separate experiments. **c**, Cryo-EM maps of FhuA-pb5 shown within the OM plane. FhuA is coloured green and pb5 magenta. LPS or detergent density is in cyan. **d**, Cartoon model of FhuA-pb5 coloured in rainbow representations (blue; N-terminus). **e**, Superposition of AlphaFold-predicted pb5 (light blue) and pb5 bound to FhuA (rainbow). Note the absence of the putative pb4-interacting domain of pb5 within the complex. **f**, Cartoon viewed from the OM plane, highlighting the interacting loops from pb5 (L1-L4) and FhuA (EL7 and EL8). **g**, Superposition of free FhuA (crystal structure; PDB ID 1BY3) and FhuA within the FhuA-pb5 complex.

The receptor for phage T5 was identified as the outer membrane (OM) TonB-dependent transporter (TBDT) FhuA almost fifty years ago^16,17^. Purified FhuA and pb5 form a highly stable complex^18,19^ that was characterised at low resolution by small angle neutron scattering and negative stain electron microscopy^20^. Taking into account the low resolution, no major conformational changes were observed upon complex formation, contrasting with multi-copy RBPs that bind to surface polysaccharides with low affinities ^21^. Addition of purified pb5 to *E. coli* cells blocks subsequent T5 infection and affects other processes that depend on functional FhuA, such as ferrichrome import ^19^. Similar phenotypes are observed upon expression of the small phage lipoprotein Llp (Uniprot Q38162), which is adjacent and downstream to pb5 on the T5 genome, establishing it as the phage T5 SE protein ^22-24^. Llp, also termed lytic conversion protein^22^, has low sequence similarity to the family of small Cor lipoproteins that infect T1 and related coliphages^25^. An *in vitro* study placed Llp on the outside of the cell^24^, but *in vivo* work suggested Llp to be periplasmic^23^. A model was proposed in which Llp, by binding to FhuA, would cause allosteric conformational changes in the pb5 binding site on the extracellular surface, thereby preventing T5 binding^23^.

To elucidate the mechanism of SE via potential TBDT structure modulation, we report here the cryo-EM structure of the FhuA-pb5 complex and the X-ray crystal structure of the FhuA-Llp complex. The FhuA-Pb5 structure shows that pb5 is an elongated molecule with one end inserted into the extracellular lumen of the FhuA barrel and with its long axis approximately perpendicular to the OM plane. All extracellular FhuA loops except EL1-EL3 contact pb5, providing a qualitative explanation for the high stability of the interaction. Free FhuA and FhuA within the complex are virtually identical. The domain of pb5 that interacts with pb4 is poorly ordered, providing a possible mechanism for transmitting a conformational change from pb5 to the rest of the tail. The FhuA-Llp structure shows that Llp is indeed bound to the periplasmic face of FhuA, making extensive interactions with the FhuA plug. Strikingly, the conformation of the plug within the complex is non-native, suggesting that Llp has bound to a FhuA intermediate state during TonB-dependent transport. On the extracellular side, FhuA loops EL7 and EL8 have undergone large conformational changes to fold inwards and completely block access to the plug domain. EL7 and EL8 would clash with pb5, providing an explanation for small lipoprotein-mediated SE via modulation of receptor structure.

## Results

To explore the pb5/FhuA/Llp interactions *in vitro*, individual components were expressed in *E. coli* and purified via IMAC and SEC. While pb5 is a soluble protein and does not stably associate with detergent micelles (unlike Llp and FhuA), the addition of detergent to *E. coli* lysates improved pb5 yield and behaviour on SEC, and pb5 was therefore purified in the presence of detergent. For FhuA-Llp, we observed that complex formation via addition of purified Llp to FhuA is very slow (Extended Data Fig. 1), suggesting that the FhuA conformation to which Llp binds is poorly accessible or sparsely populated *in vitro*. To obtain the FhuA-Llp complex, we co-overexpressed FhuA and Llp on different plasmids in *E. coli* and purified the *in vivo*-assembled complex via IMAC and SEC. Depending on the particular preparation, we could obtain a roughly equimolar complex that is very stable during gel filtration (Extended Data Fig. 1).

We next analysed the interaction of the proteins via size exclusion chromatography (SEC). Mixing of equimolar amounts of pb5 and FhuA followed by short incubation (5 min) results in one peak on SEC containing both components (Fig. 1b) and no trace of free pb5, indicating formation of a very stable FhuA-pb5 complex. By contrast, mixing equimolar amounts of pb5 and FhuA-Llp yields two well-separated peaks on SEC, with no pb5 co-eluting with FhuA-Llp. It should be noted that pb5 elutes much later than expected on SEC, and the FhuA-pb5 complex runs only slightly faster than FhuA alone (Fig. 1b). These data show that the prevention of pb5 binding to the phage T5 receptor FhuA by the phage lipoprotein Llp can be reconstituted *in vitro* with purified components, and no other factors are required.

Since pb5 bound to FhuA is more stable at high concentrations (> 0.5 mg/ml) than pb5 in isolation, the FhuA-Pb5 complex was obtained in milligram amounts by mixing pre-purified FhuA with *E. coli* cell lysates expressing pb5, followed by IMAC and gel filtration in the presence of detergent (Fig. 1b). Crystallisation trials yielded crystals diffracting anisotropically and only to modest resolutions (∼4 Å), and the phase problem could not be solved by molecular replacement (MR) with FhuA (PDB ID 1BY3)^26^ and AlphaFold2-predicted pb5 as search models ^27,28^. We succeeded in solving the FhuA-pb5 structure via cryo-EM, using complex purified in decyl-β-maltopyranoside (DM) at 3.1 Å resolution (Extended Data Fig. 2 and Supplementary Table 1). The ∼150 kDa complex is ∼150 Å high and has a largest width of ∼65 Å at the base of the FhuA barrel (Fig. 1c). The LPS molecule that is present in FhuA X-ray crystal structures^29^ is clearly visible in the cryo-EM map. Consistent with the AlphaFold2 prediction of isolated pb5, FhuA-bound pb5 has an oblong shape with a large central β-sheet (Fig. 1d). Like many phage proteins, pb5 is not similar to any other protein; a DALI analysis^30^ identifies PDB ID 2GSY (polyprotein) as the closest structural homolog with a Z-score of 5.3, but with only 99 aligned residues (Cα r.m.s.d. 3.5 Å; Extended Data Fig. 3). The central region of Pb5 is virtually identical to that in the AlphaFold prediction, while the part that interacts with FhuA shows large differences (overall r.m.s.d. 1.7 Å for 529 out of 640 Cα atoms; Fig. 1e and Extended Data Fig. 4). Interestingly, the region at the other end of pb5 that most likely interacts with pb4 is poorly ordered but present in the cryo-EM density, and only visible at low contours (Extended Data Fig. 4). The predicted structure of this region shows pseudo 3-fold symmetry, suggesting that pb4 may be trimeric. Overall, 529 out of 640 pb5 residues could be modelled. Pb5 inserts four loops (numbered L1-L4 from the N-terminus) into the extracellular lumen of FhuA (Fig. 1f). The involvement of L4 (residues 570-590) corrects the previous assigment of the N-terminal half of pb5 as the FhuA-interacting domain^31^. The conformations of the binding loops, as well as those of the other loops of pb5 that interact with FhuA, are predicted with low confidence by AlphaFold2, so it is unclear whether their very different conformations within the complex are caused by the FhuA interaction (Fig. 1e and Extended Data Fig. 4). Pb5 residues Gln115 in the tip of L1 and Phe170 in the tip of L2 contact Phe115 and Tyr116 in the FhuA plug (residue numbering for the mature part of FhuA), but it is clear that the insertion of pb5 does not affect the position and conformation of the plug (Fig. 1f). In fact, the entire FhuA structure remains remarkably similar upon pb5 binding, as judged from a comparison with the FhuA crystal structure (Cα r.m.s.d 0.8 Å with 1BY3; Fig. 1g). With the exception of EL1-EL3, all FhuA loops contact pb5, resulting in a large interface area of ∼2140 Å^2^ as analysed via PISA^32^. There are 27 inter-molecular hydrogen bonds, with most of them between EL4-L2 (8), EL5-L2 (8) and EL8-L1-3 (7) (Extended Data Fig. 5a). Our structure supports the data from a study on the effect of systematic FhuA loop deletions on T5 infection, where deletion of any individual loop, with the exception of EL8, had only a modest effect on T5 sensitivity^33^. Pb5 completely fills the extracellular lumen of FhuA and occludes the ferrichrome binding site (Extended Data Fig. 6), explaining why addition of purified pb5 to *E. coli* inhibits growth under iron-starved conditions^19^. Interestingly, pb5 binding does not cause conformational changes in the Ton box, and the structure of the plug in FhuA-pb5 is identical to that in apo FhuA, despite the fact that ferrichrome and pb5 both contact Phe115 and Tyr116 of the plug (Extended Data Fig. 6). Thus, phage T5 is not recognised as a ligand by FhuA.

Having characterised the FhuA-Pb5 interaction at high resolution, we next focused on solving the structure of the FhuA-Llp complex. Depending on the particular preparation, we could obtain a roughly equimolar complex that is stable during gel filtration. Due to the relatively small size of FhuA-Llp (∼90 kDa), we utilised X-ray crystallography for this part of the project. Extensive screening yielded one crystal form that contained both components (Extended Data Fig. 1) and had useful, anisotropic diffraction to ∼3.4 Å resolution. MR with FhuA resulted in maps with unaccounted density on the periplasmic face of FhuA, but of insufficient quality for model building. Adding an AlphaFold2-predicted Llp model to the MR search gave a solution that allowed building and refinement of the complete FhuA-Llp complex (Fig. 2a and Supplementary Table 2). The structure shows that the prediction of Llp, while providing valuable phasing information, is inaccurate overall (Cα r.m.s.d 2.8 Å for 28 aligned atoms out of 61; Fig. 2b). In addition, the AlphaFold2-predicted complex erroneously places Llp on the extracellular side (Extended Data Fig. 7), at a position that is impossible to reconcile with the lipid anchor at the N-terminus of Llp. The other two cysteines in Llp (Cys11 and Cy58) are disulphide-bonded, constraining both termini of the protein.

**Figure 2.**
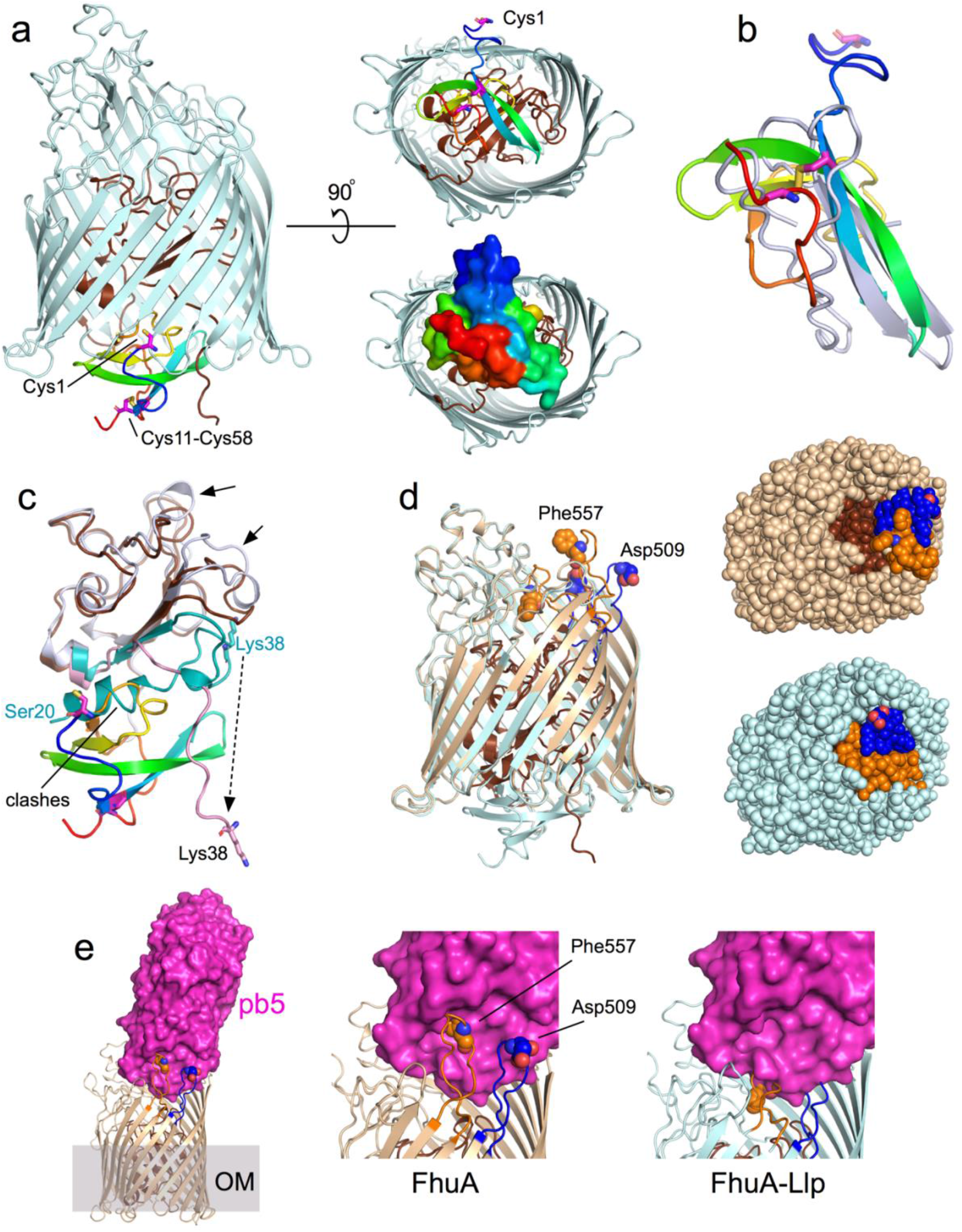
Llp binding to FhuA generates allosteric conformational changes in the transporter that abolish pb5 binding. **a**, Cartoon viewed from the OM plane showing Llp bound to the periplasmic face of FhuA (light blue; plug, brown). Llp is coloured in rainbow representation (N-terminus blue) and the cysteines are shown by magenta stick models and labeled. The right panels show views from the periplasmic space, with a surface model of Llp shown at the bottom. **b**, Superposition of AlphaFold2-predicted Llp (blue-gray) and Llp bound to FhuA (rainbow). **c**, Llp binds to a non-native state of the FhuA plug. The plug of free FhuA is blue-gray, with residues Ser20-Glu57 coloured teal. The plug of Llp-bound FhuA is brown, with residues Lys38-Glu57 pink. Residue Lys38 is shown for both structures as sticks. “Clashes” indicate the region of the free FhuA plug that would overlap with Llp. Arrows show movements of plug segments on the extracellular side of the plug. View direction is roughly as in left panel (**a**), with the periplasmic space at the bottom. **d**, Extracellular changes in FhuA caused by Llp binding. Superposition of free FhuA (wheat) and FhuA-Llp, with plug domains shown in brown. FhuA loops EL7 (Glu501-Ser516) and EL8 (Thr546-Glu564) are coloured blue and orange, respectively, with residues Asp509 (EL7) and Phe557 (EL8) shown as space-filling models. The right panels show space-filling models from the outside of the cell, demonstrating the occlusion of the plug (brown) by EL7 and EL8 movement. **e**, Cartoon of free FhuA with the pb5 surface superimposed. The central and right panels show close ups of the EL7/EL8 interface with pb5 for free FhuA and FhuA-Llp, respectively.

Llp is bound to the periplasmic face of FhuA and makes extensive interactions with both the plug and the barrel. The total interface area is 1640 Å^2^, with 14 hydrogen bonds and four salt bridges (Extended Data Fig. 5). Many of the interactions occur between Llp and the visible N-terminal ∼20 residues of the plug, comprising Lys38-Glu57. Interestingly, those plug residues have a very different conformation in free FhuA. In addition, density up to Ser20 is visible in free FhuA, which includes the N-terminal plug helix. Pairwise backbone differences between residues visible in both structures are as much as 26 Å for Lys38 (Fig. 2c). Strikingly, several Llp residues (Ile39-Trp46) occupy space where the N-terminal plug helix of free FhuA would be, indicating that Llp has bound to a non-native conformation of FhuA (Fig. 2c and Extended Data Fig. 8). Structural changes in the rest of the plug are less dramatic, but nevertheless still include backbone shifts of 3-4 Å downwards, towards the periplasmic space.

On the extracellular side, the conformational changes in FhuA-Llp relative to free FhuA are dramatic but confined to just two loops, EL7 and EL8 (Fig. 2d). Both loops fold inwards to completely occlude the plug domain in Llp-bound FhuA. Backbone shifts for residues located at the loop tips (Asp509 in EL7 and Phe557 in EL8) are ∼15 Å (Fig. 2d). To investigate the possibility that the conformations of EL7 and EL8 are induced by the crystallisation process we performed unbiased molecular dynamics (MD) simulations on the FhuA-Llp system. Three independent simulations show that the loop conformations are stable on the timescales of the simulations (2 μs; Extended Data Fig. 9), suggesting they have been induced by Llp binding to FhuA and not by the crystal lattice (Extended Data Fig. 10). Llp is mobile but remains bound to FhuA, suggesting that the interactions of Llp with the plug dominate those with the FhuA barrel. Importantly, the effect of the EL7 and EL8 movements is that they will prevent the high-affinity binding of pb5 to FhuA because of extensive clashes with the FhuA binding loops of pb5 (Fig. 2e), explaining our *in vitro* data with purified components and *in vivo* data from the literature (Fig. 1b)^19,22-24^. Moreover, the Llp-induced loop positions will also prevent albomycin binding (Extended Data Fig. 6)^34^, explaining why cells co-expressing FhuA and Llp become resistant towards this antibiotic^23^. Llp binding to FhuA also generates resistance of *E. coli* towards phage Φ80 and colicin M^23^, suggesting that FhuA loops EL7 and EL8 are also involved in the binding of those phages. Interestingly, the deletion of EL8 has the biggest effect on phage T5 sensitivity of all FhuA loops^33^, potentially explaining why this particular loop, together with EL7, undergoes a conformational change on Llp binding.

## Discussion

An early study^23^ investigated the effects of Llp on FhuA-dependent processes such as phage T5 sensitivity for wild type and mutant transporters. Unfortunately, this study, like others performed around the same time, predated the crystal structures of FhuA and assumed a wrong topology model. It would in any case be hard to rationalise the effect of most FhuA mutations without detailed structural and functional analyses of their effect on FhuA structure and FhuA-Llp/FhuA-pb5 interactions. A notable exception is the I9P mutation made for the Ton box of FhuA, which is the only mutant to revert the inactivation of FhuA by Llp, *i*.*e*., FhuA I9P cells remain sensitive towards T5 infection in the presence of Llp^23^. The proline substitution disrupts the β-strand structure of the Ton box and likely abolishes the interaction with TonB, as suggested by the fact that the cells become resistant to albomycin^23^. Since neither Llp nor pb5 interacts with the Ton box, the FhuA I9P data support our hypothesis that Llp binds to an intermediate state of FhuA that is populated during the TonB-dependent transport cycle (Fig. 3). The Ton box in FhuA-Llp (residues Asp7-Ala14) is not visible presumably owing to flexibility, but given the exposed location of Lys38 (Fig. 2c) it seems reasonable to assume the Ton box is accessible to TonB in FhuA-Llp. We propose that Llp has jammed the plug of FhuA, and that TonB pulling does not remove the phage lipoprotein from the transporter. In addition, due to the presence of Llp, the periplasmic face of the plug cannot interact with TonB^35^.

**Figure 3.**
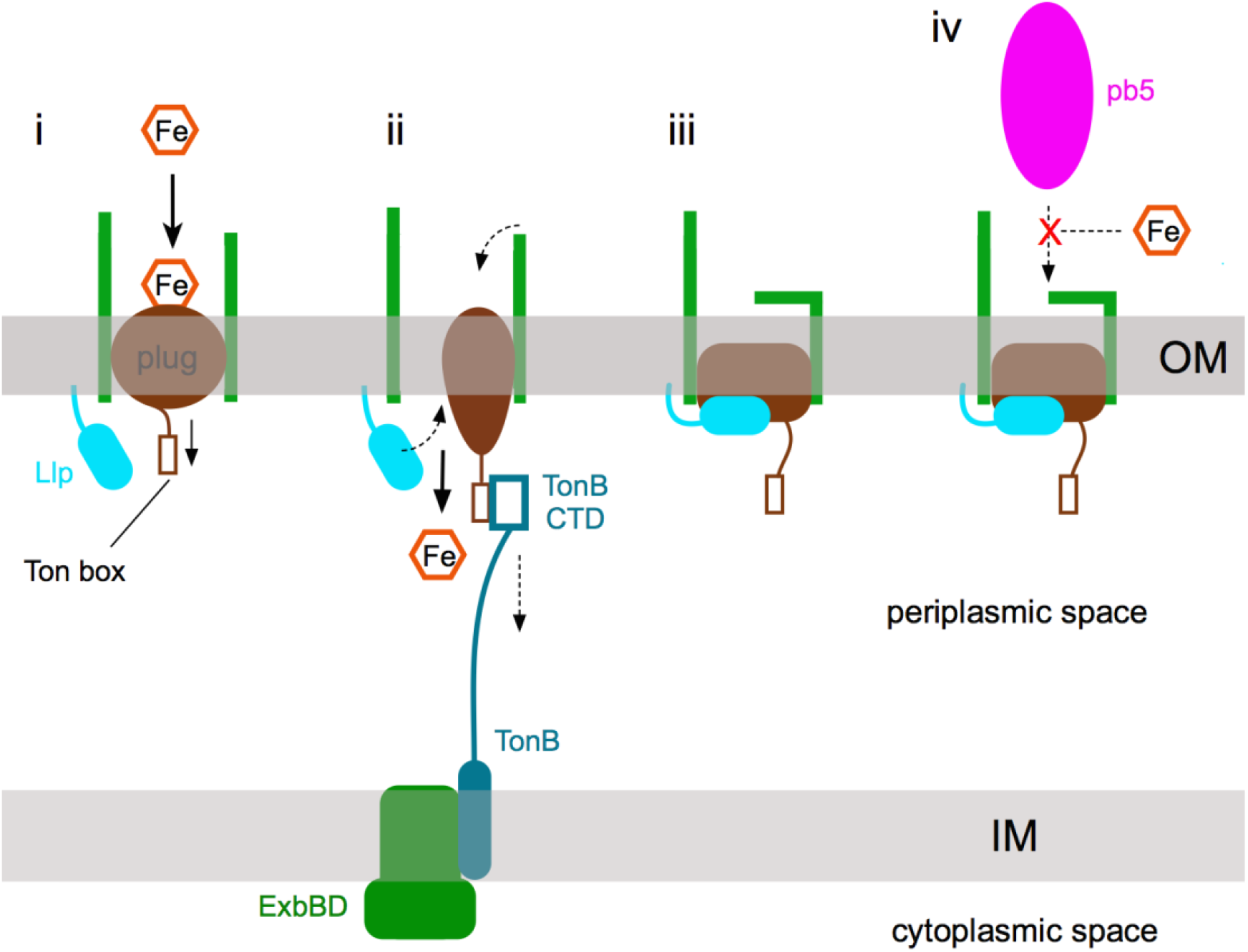
Schematic model for lipoprotein-mediated SE via TonB-dependent transporters. (**i**) Substrate (in this case iron-siderophore) binding to the TBDT (green) causes allosteric changes in the plug (brown) that expose the Ton box to the periplasmic space. (**ii**) The C-terminal domain (CTD) of TonB binds the Ton box and, by mechanical force, causes a conformational change in the plug that allows substrate passage into the periplasmic space. (**iii**) During the transport cycle, the phage SE lipoprotein (Llp; cyan) binds to the plug, causing allosteric changes that generate conformational changes in one or more extracellular loops, which abolish binding of phage RBPs, here represented by pb5 (**iv**). The SE lipoprotein jams the TBDT in a transport-incompetent state despite the Ton box being accessible.

Our structures show that the phage lipoprotein Llp exploits the known allostery in the plug of FhuA in the opposite direction as during ferrichrome import, *i*.*e*. from the periplasmic side to the extracellular side. How exactly Llp binding causes the conformational changes in EL7 and EL8 remains difficult to answer. The plug loops indicated by the arrows in Fig. 2c contact the base of EL7 and EL8, and their downward movement in FhuA-Llp creates space that may allow EL7 and EL8 to fold inwards. The tips of EL7 and EL8 make many interactions with barrel and loop residues; in particular, the side chains of Phe557 and Phe558 in EL8 interact with an extensive aromatic patch, stabilising the inward-folded conformation of EL8.

Llp has low pairwise sequence identity (∼20%, including the intramolecular disulphide bond) to members of the Cor small lipoprotein family^25^ that mediate SE in *e*.*g*. phage T1 and Φ80. These phages, like T5, use FhuA as terminal receptor and their Cor proteins likely function in a similar way as Llp. Since the “forward” allostery in TBDTs leads to Ton box exposure in the periplasmic space regardless of the identity of transporter and substrate, it seems plausible that the lipoprotein-mediated SE we describe for FhuA is utilised by many other TBDTs. Indeed, a recent study suggests that TBDT-targeting SE lipoproteins are common in coliphages^36^.

Considering phage diversity and the notion that SE seems advantageous for the phage, it is likely that there are many ways by which SE can be achieved. As an example, a recent study showed that SE mediated by the phage T4 protein Spackle is caused by the binding of Spackle to the lysozyme domain of the T4 tail spike protein gp5, inhibiting its activity^37^. While little is currently known, a common feature of SE proteins seems to be their small size, and it is conceivable that a considerable fraction of phage proteins of unknown function, many of which are small (< 10 kDa), may be involved in SE. Regarding the SE mechanism we have discovered here, in which an OM lipoprotein modulates the structure of a protein receptor, an intriguing question is whether this also occurs in phages that have non-TBDTs as receptors, and how the cognate RBPs are blocked. Mafei et al. recently characterised the host specificity of a new library of coliphages experimentally^36^, showing that while most Siphophages infecting *E. coli* use a TBDT as terminal receptor (*e*.*g*. FhuA, FepA, BtuB), a substantial number of phages target LamB, TolC or LptD, none of which are TBDTs. The putative RBPs and cognate SE proteins of most of these phages can be identified based on their location directly downstream of the *gpJ* locus, which encodes a tail tip protein homologous to the tail J protein of phage lambda. Inspection of the downstream region of *gpJ* reveals that phages that target LamB (*e*.*g*. Bas23) or LptD (*e*.*g*. Bas18 and phage RTP) likely do have SE lipoproteins^36^. Since neither LamB nor LptD are known to have allostery, the question remains how those lipoproteins could block the phage receptor. *E. coli* can flip “standard” lipoproteins such as Braun’s lipoprotein to the cell surface^38^, so SE lipoproteins blocking *e*.*g*. LptD may be located on the extracellular side of the OM rather than being periplasmic.

A major question is how Siphophage DNA is transferred into the host cell. Our structural data for FhuA-pb5 provide the most direct evidence to date for the view that DNA transfer does not occur via the FhuA channel, as originally proposed based on single channel electrophysiology^39,40^. Contrasting with an earlier proposal based on cryo-ET^41^, our structure shows that pb5 is bound in the center of FhuA, and not at the periphery, confirming previous negative stain electron microscopy^20^. Combined with data placing pb5 at the very tip of the central tail fiber (Fig. 1a), and the loss of tail tips when detergent-purifed FhuA is incubated with T5 (ref. 13), we propose that the disorder observed in the pb4-interacting part of pb5 (Extended Data Fig. 4) is caused by FhuA binding and leads to the dissociation of the tail tip. This would then result in the TMP (pb2), located within the pb6 tail tube, becoming available for cell envelope perforation. The mechanism for this process remains obscure, but it is worth noting that peptidoglycan-degrading enzymes (endolysins), when fused to pb5, can lyse *E. coli* cells in a FhuA-dependent manner^42^, suggesting that close and prolonged proximity to the surface is crucial for compromising OM integrity. For phage T5, OM proximity after tail tip loss would be maintained by the lateral fibers (pb1), which have LPS as primary receptor. We speculate that the “two-tier” interaction with OM receptors (LPS followed by FhuA) exists because the pb1 interactions with LPS are not sufficient to generate the conformational changes in the tail tip that are required for DNA transfer into the host cell.

## Acknowledgements

We thank Lone Brøndsted (University of Copenhagen) for the gift of pb5 DNA and Javier-Abellon-Ruiz for generating the Bl21 (DE3) Δ*cyo* expression strain. We would like to acknowledge the Diamond Light Source for crystallography beam line access (proposal mx-24948) and i04 beamline support, as well as eBIC for rapid access time (proposal BI31384). SLM acknowledges the support of the Federation of European Biochemical Societies (FEBS) through a long-term fellowship. The research of BvdB is supported by a Wellcome Trust Investigator award (214222/Z/18/Z), providing salary support for AS. SK is funded by the EPSRC via HECBioSim.

## Author contributions

BvdB conceived the project, performed experiments, and wrote the paper. AS, together with BvdB, determined the cryo-EM structure of FhuA-pb5 and assisted with writing the paper. AB collected X-ray crystallography and cryo-EM data. SM performed MD simulations, supervised by SK.

## Methods

### Cloning, protein expression and purification

The mature part of *E. coli* FhuA (starting with residue 34) was amplified from *E. coli* gDNA and cloned into a modified version of pET9 via ligation-independent cloning (LIC), generating a construct with the PelB signal sequence followed by a His10 tag and tobacco etch virus (TEV) cleavage site. The gene coding for pb5 was amplified by PCR, digested with NcoI and XhoI, and ligated into pET28-b restricted with the same enzymes, adding the sequence “LE” followed by a His6 tag to the C-terminus of the protein. For Llp, a codon-optimised gene (Eurofins Genomics) for expression in *E. coli* was digested with EcoRI and XbaI and ligated into the arabinose-inducible vector pB22 restricted with the same enzymes. A C-terminal His6 tag was included in the synthetic gene.

FhuA expression was performed in the Bl21 (DE3) Δ*cyo* strain that has a clean deletion of the *cyoAB* ubiquinol oxidase genes, abolishing contamination of IMAC samples with the abundant Cyo complex, and removing the need for selective removal of IM proteins via *e*.*g*. sarkosyl pre-extraction steps. Cells were grown in LB at 37 °C and 180 rpm to an OD_600_ 0.2-0.4 (50 μg/ml kanamycin), and placed in the cold room for 20-30 min prior to induction with 0.2 mM IPTG. The cells were grown for another 18-20 hrs at 18 °C and 150 rpm. Llp was expressed either in isolation or together with FhuA in Bl21 (DE3) Δ*cyo* as described above. Growth of cells at 37 °C gave substantially higher yields of Llp compared to growth at lower temperatures. For co-expression, Bl21 (DE3) Δ*cyo* cells were transformed with pET9 and pB22 plasmids via electroporation of freshly made electrocompetent cells and plated out on LB plates with kanamycin (50 μg/ml) and ampicillin (100 μg/ml). For liquid cultures, 35-40 μg/ml kanamycin was used to avoid excessive lag phases and cells were induced with 0.2 mM IPTG and 0.1% (w/v) (L)-arabinose. Expression of pb5 was performed in Bl21 (DE3) at 18 °C as described above for FhuA. Final OD_600_ values were typically 2-3.

Cells were harvested by centrifugation for 20 min at 4,200 rpm (JS 4.2 rotor) in a Beckman J6-HC centrifuge. Cell pellets were processed either directly or frozen at -20 °C. Cells were resuspended in TSB buffer (20 mM Tris, 300 mM NaCl pH 7.8), homogenised by douncing, and lysed in a cell disrupter (Constant Systems 0.75 kW model) at 20-23,000 psi (1 pass) in the presence of DNAse. Typically 120 ml of buffer was used for 2-4 liters of cells. The lysed cells were centrifuged at 42,000 rpm (45Ti rotor; Beckman Optima XE-90) for 50 min and either the supernatant (in the case of pb5) or the total membrane pellet (for FhuA and/or Llp) was collected. Membranes containing overexpressed FhuA, Llp, or FhuA-Llp were extracted with either 1.5% LDAO or 2.5% Elugent (typical volume ∼60 ml for membranes from 2 l culture) by douncing, folllowed by stirring for 2 hrs at 4 °C, or in some cases, overnight. No difference was observed in extraction efficiency or downstream purification, and no protease inhibitors were added. Following extraction, the suspension was centrifuged at 42,000 rpm for 30 min and the clarified extract was subjected to IMAC using Ni-charged Chelating Sepharose (Cytiva) equilibrated in TSB + 0.15% LDAO. Following loading, the column was washed with ∼20 column volumes buffer with 30 mm imidazole, and eluted with ∼3 column volumes buffer with 200 mM imidazole. IMAC elutions were immediately concentrated by centrifugal filtration (Amicon Ultra-15; MWCO 50 kDa for FhuA and 30 kDa for Llp), and subjected to size exclusion chromatography (SEC) on Superdex-200 16/10 in (typically) 10 mM Hepes, 100 mM NaCl, 0.05% LDAO pH 7.5, and appropriate peak fractions were collected. In the case of FhuA and Llp co-overexpression, preparations from cells grown at 37 °C showed, when analysed via SEC, in addition to a FhuA-Llp a peak for free Llp, indicating that Llp is present in excess to FhuA in the *E. coli* OM. For FhuA-Llp crystallisation, detergent exchange was performed by a second SEC column in which the LDAO was replaced by either 0.4% C_8_E_4_ or 0.25% DDAO. Protein was concentrated to 10-15 mg/ml, flash-frozen in LN2 and stored at -80 °C. The FhuA-Llp preparation that gave the diffracting crystals was treated with TEV protease following SEC, using TEV buffer (50 mM Tris, 0.5 mM EDTA, 0.2 mM TCEP pH 8) containing 0.05% DDM. A ratio of TEV:FhuA-LLp of ∼10 (w/w) was used and the incubation was done at 4 °C for 16 hrs. Following cleavage, TEV was separated from FhuA-Llp via SEC in 0.4% C_8_E_4_, and protein was concentrated to 10-15 mg/ml, aliquoted, flash-frozen in LN2 and stored at -80 °C.

Following ultracentrifugation, the supernatant of pb5-expressing cells was loaded on IMAC and processed as above in the absence of any detergent. However, analysis on SEC (10 mM Hepes, 100 mM NaCl pH 7.5) showed that pb5 eluted as a broad peak. We subsequently observed that adding detergent to the pb5 sample improved the behaviour on SEC. We therefore added 0.1% LDAO to the supernatant followed by pre-purified FhuA, ensuring that pb5 was in at least 2-fold excess over FhuA. Following a short (15 min) incubation, the supernatant was loaded on IMAC and processed as for free FhuA. The IMAC elution was concentrated (50 kDa MWCO cutoff) and loaded on a Superdex-200 16/60 column equilibrated in 0.05% LDAO as described above. Peaks corresponding to FhuA-pb5 and free pb5 were collected, concentrated to ∼8-10 mg/ml and 0.5 mg/ml respectively, and flash-frozen in LN2. To improve the stability of free pb5, 10% glycerol was added prior to flash-freezing.

For *in vitro* interaction studies, 1 nmole of each protein was mixed and incubated at room temperature for ∼15 min, and ran on a Superdex-200 Increase 10/300 GL equilibrated in 10 mM Hepes/100 mM NaCl, 0.05% LDAO pH 7.5 (injection volume ∼0.4 ml, flow rate 0.5 ml/min).

### Cryo-EM data acquisition for FhuA-pb5 and data processing

Purified FhuA-pb5 complex (3.5 μl) at 8 mg/ml was applied to glow discharged (15 mA, 40 s) Quantifoil 1.2/1.3 300 mesh holey carbon grids. The grids were immediately blotted and plunge-frozen in liquid ethane using a Vitrobot Mark IV (ThermoFisher Scientific) device operating at 4°C and ∼100% humidity. Data were collected on a FEI Titan Krios microscope operating at 300 kV using a Falcon 4 direct electron detector with a Selectris imaging filter (10 eV slit width) (ThermoFisher Scientific) at the Astbury Biostructure Laboratory (Supplementary Table 1). A total of 8,387 movies were recorded in counting mode at 165,000 magnification, corresponding to a pixel size of 0.71 Å.

All image processing was done in cryoSPARC v.3.3.2 (refs. 43, 44). Movies were motion corrected using patch motion correction, and CTF parameters were estimated using patch CTF estimation. 6,566 micrographs remained after discarding average intensity, defocus and ice thickness outliers. Initially ∼2,000 particles were picked manually, subjected to 2D classification and the resulting 2D classes were used for template-based picking. 2,054,732 particles were extracted in 360 pixel boxes. 2D classification was used to discard bad particles, followed by generation of an ab initio 3D model using a stochastic gradient descent approach with 2 classes. Particles from the best looking class were subjected to non-uniform refinement, resulting in a 3.57 Å reconstruction from 74,313 particles. The particles were re-extracted with a box size of 440 pixels and the final set of 71,476 particles was used in non-uniform refinement with CTF parameter (beam tilt and trefoil) and per-particle defocus estimation. The final reconstruction had a global resolution of 3.1 Å, with the protein core regions reaching 2.7 Å as assessed by local resolution estimation (Extended Data Fig. 2). The FhuA crystal structure and the AlphaFold2 model of pb5 were rigid-body fit in the map via Phenix DockinMap^45^, and the model was built via several cycles of manual building in COOT^46^ and real-space refinement within Phenix. The final model refinement statistics are listed in Supplementary Table 1.

### Crystallisation and structure determination of FhuA-Llp

Preparations of FhuA-Llp purified in either DDAO or C_8_E_4_ were subjected to initial crystallisation screening via sitting drop vapour diffusion, by mixing 200 nl of protein with 200 or 150 nl well solution via a Mosquito crystallisation robot (TTP Labtech) at 20 °C. Several commercial screens were typically used (MemGold, MemGold2, MemTrans and MemChannel; Molecular Dimensions). Several hits were obtained in C_8_E_4_, but only one of these (16% PEG 4K, 0.4 M (NH_4_)_2_SO_4_, 0.1 M NaAc pH 4.5) diffracted occasionally to beyond 5 Å resolution. The size of the initial crystals was increased manually via hanging drop vapour diffusion with larger drops (typically 1-1.5 μl), and further fine screening around the initial hit condition led to the collection of a best dataset with moderately anisotropic diffraction to ∼3.4 Å resolution. Molecular replacement via Phaser^47^ within Phenix^45^ using FhuA as a search model (PDB 1BY3) gave a definite solution for FhuA and density on the periplasmic side of the plug, but of insufficient quality for model building. Addition of an AlphaFold2-predicted model for Llp provided a solution that allowed building of the complete model for Llp via several cycles of model building in COOT^46^ and refinement via Phenix. During refinement, automatically assigned TLS was used and X-ray/ADP weights were optimised to keep the refinement stable, resulting in tight r.m.s.d. values for the bond lengths and angles. Moreover, using data processed via AUTOSOL/STARANISO^48,49^ provided the best refinement results. Data collection and refinement statistics are summarised in Supplementary Table 2. Interaction surfaces were analysed via the PISA webserver at https://www.ebi.ac.uk/msd-srv/prot_int/pistart.html.

### Molecular dynamics simulations

Classical atomistic molecular dynamics (MD) simulations were initiated from the resolved X-ray structure of the FhuA-Llp complex. The system setup was prepared using the CHARMM-GUI webserver^50^. Lipid tails were added to the N-terminal cysteine residue of Llp, and the protein complex was embedded in an outer membrane model from *E. coli*, containing Ra-LPS without O-antigen in the outer leaflet and PVPE (1-palmitoyl 2-cis-vaccenic phosphatidylethanolamine), PVPG (1-palmitoyl 2-cis-vaccenic phosphatidylglycerol), and cardiolipin (1-palmitoyl 2-cis-vaccenic 3-palmitoyl 4-cis-vaccenic diphosphatidylglycerol) in the inner leaflet in a molar ratio of 90:5:5. The system was solvated with water containing 200 mM KCl ions, and negatively charged chemical groups of LPS were neutralized with calcium ions. Three MD simulations were performed starting from this system setup. Energy minimization was conducted for 5000 steps using the steepest descent algorithm. Six short consecutive MD simulations of 20 ns in total were performed following the CHARMM-GUI protocol employing an integration timestep of 1-2 fs, the Berendsen thermostat and barostat at a temperature of 313 K and a pressure of 1 bar^51^, and decreasing position restraints of amino acids and lipids. MD simulations of the three equilibrated systems were performed for 2 μs each using an integration timestep of 2 fs, the Verlet cutoff-scheme, and a cutoff distance for van der Waals and electrostatic interactions of 1.2 nm. The Nose-Hoover thermostat was employed at a temperature of 313 K^52,53^, and semi-isotropic pressure coupling was achieved using the Parrinello-Rahman barostat at a pressure of 1 bar^54^. Covalent bonds involving hydrogen atoms were constrained using the LINCS algorithm^55^. Electrostatic interactions were evaluated using the Particle-Mesh Ewald (PME) method^56^. MD simulations were performed using the GROMACS software package^57^ and the CHARMM36m force field^58,59^, and were visualized and analysed using the VMD software^60^.

## Data availability

The data supporting the findings of this study are available from the corresponding author upon reasonable request. Coordinates and structure factors that underly the findings of this work have been deposited in the Protein Data Bank with accession codes 8A60 (Fhua-Llp). EM structure coordinates and maps have been deposited in the Electron Microscopy Data Bank with accession codes 8A8C (FhuA-pb5) and EMD-15229.

## Extended Data

**Extended Data Figure 1.**
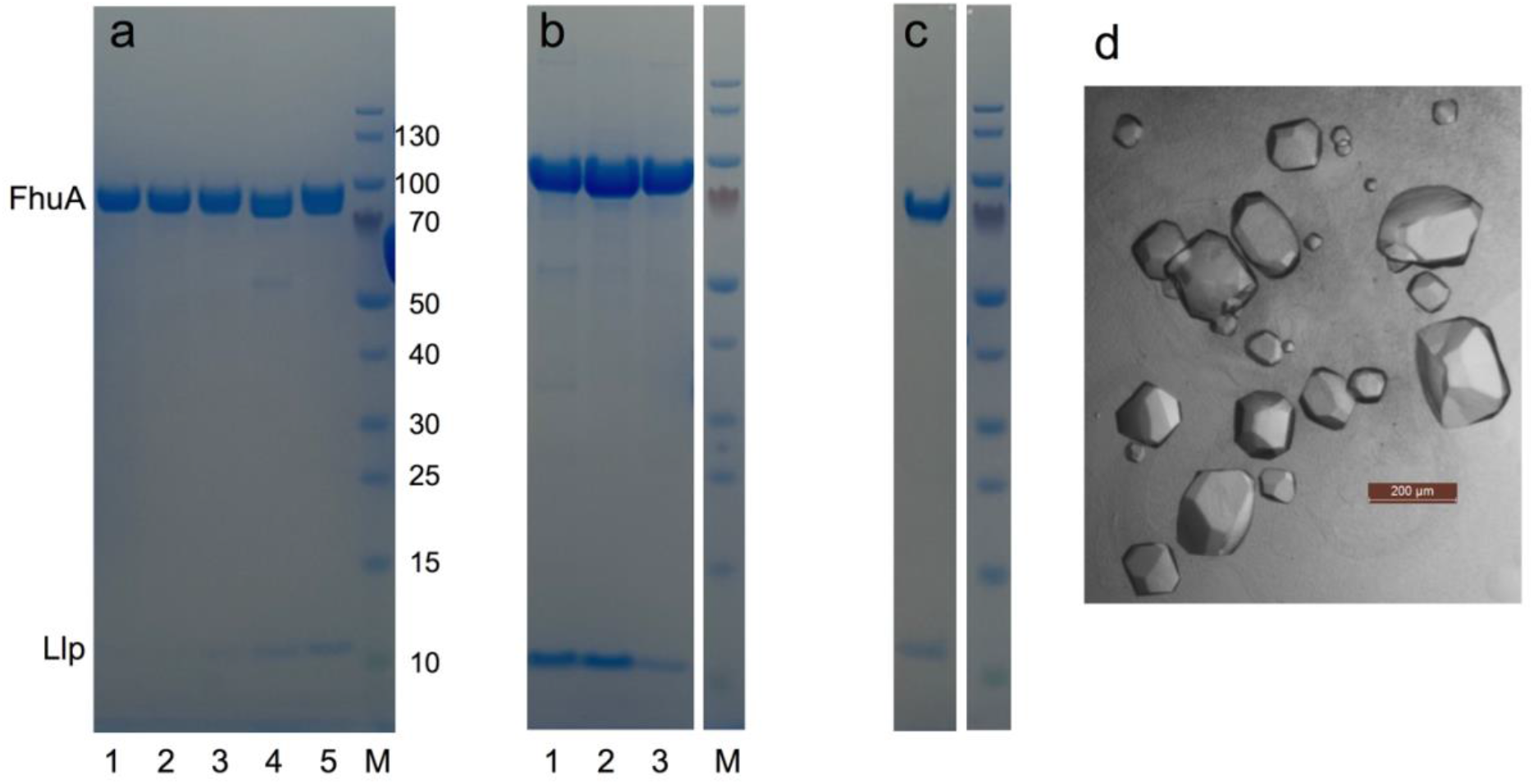
Various properties of the FhuA-Llp complex explored via SDS-PAGE. **a**, Complex formation *in vitro* is extremely slow. Purified Llp was incubated at 2-fold molar excess to FhuA and samples were analysed via SEC (Superdex-200 Increase 10/300 GL) at various time points. Lane 1, 30 min; lane 2, 60 min; lane 3, 24 hrs; lane 4, 48 hrs; lane 5, *in vivo*-generated FhuA-Llp complex purified in LDAO; M, molecular weight marker. In all cases, a sample corresponding to the FhuA peak was analysed. Due to the small size of Llp (∼7 kDa), FhuA and FhuA-Llp elute at the same volume. **b**, Variability of the FhuA:Llp ratio for different preparations, which is most likely due to different relative co-expression levels of FhuA and Llp. Lane 1, equimolar amounts of FhuA and Llp, based on individual OD_280_; lane 2, FhuA-Llp preparation 1; lane 3, FhuA-Llp preparation 2. **c**, Dissolved FhuA-Llp crystals show the presence of both components. **d**, Diffracting crystals of FhuA-Llp obtained in MemGold condition F12.

**Extended Data Figure 2.**
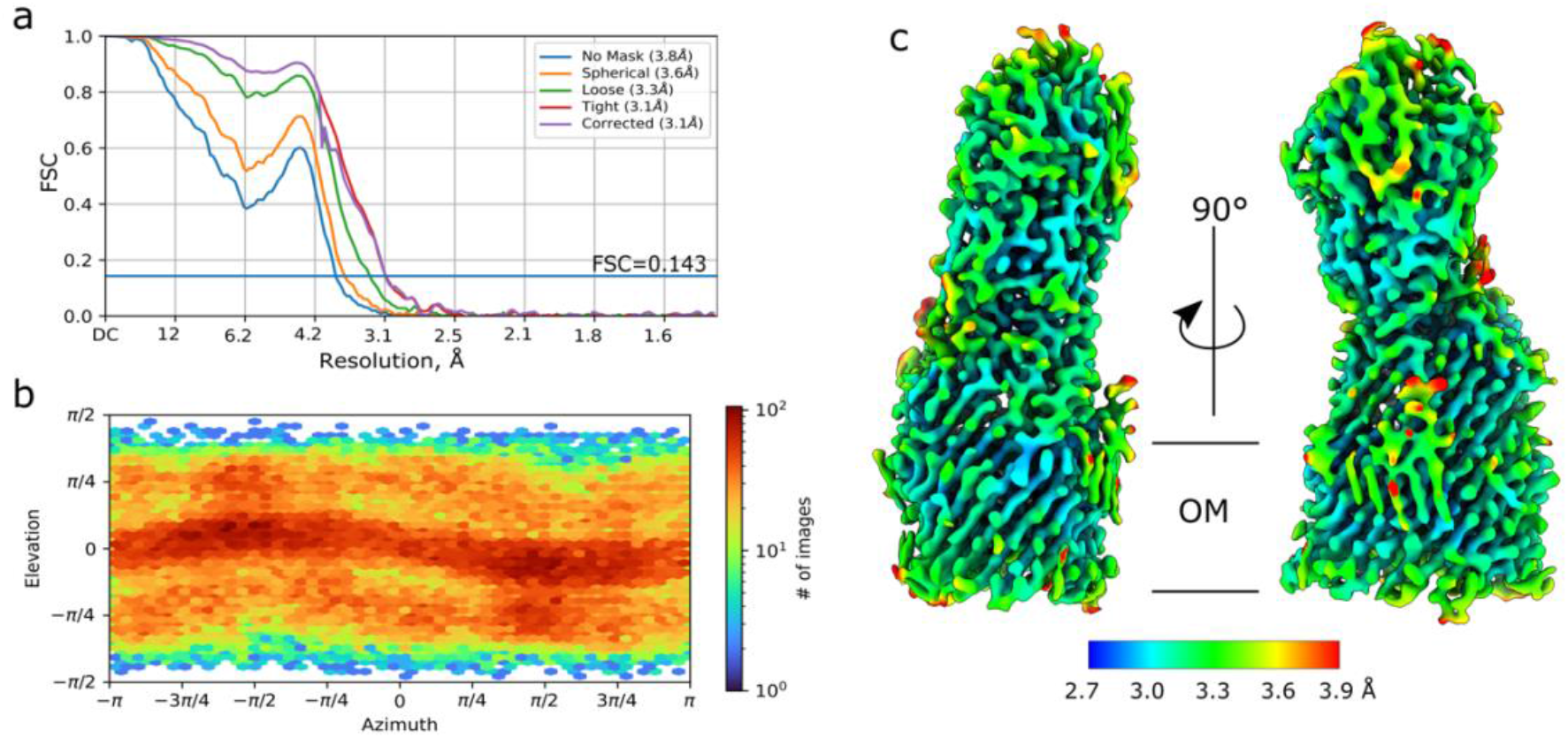
Structure determination of FhuA-pb5. **a**,**b** Gold-standard FSC curves (**a**) and viewing distribution plot (**b**). **c**, Local resolution map of FhuA-pb5.

**Extended Data Figure 3.**
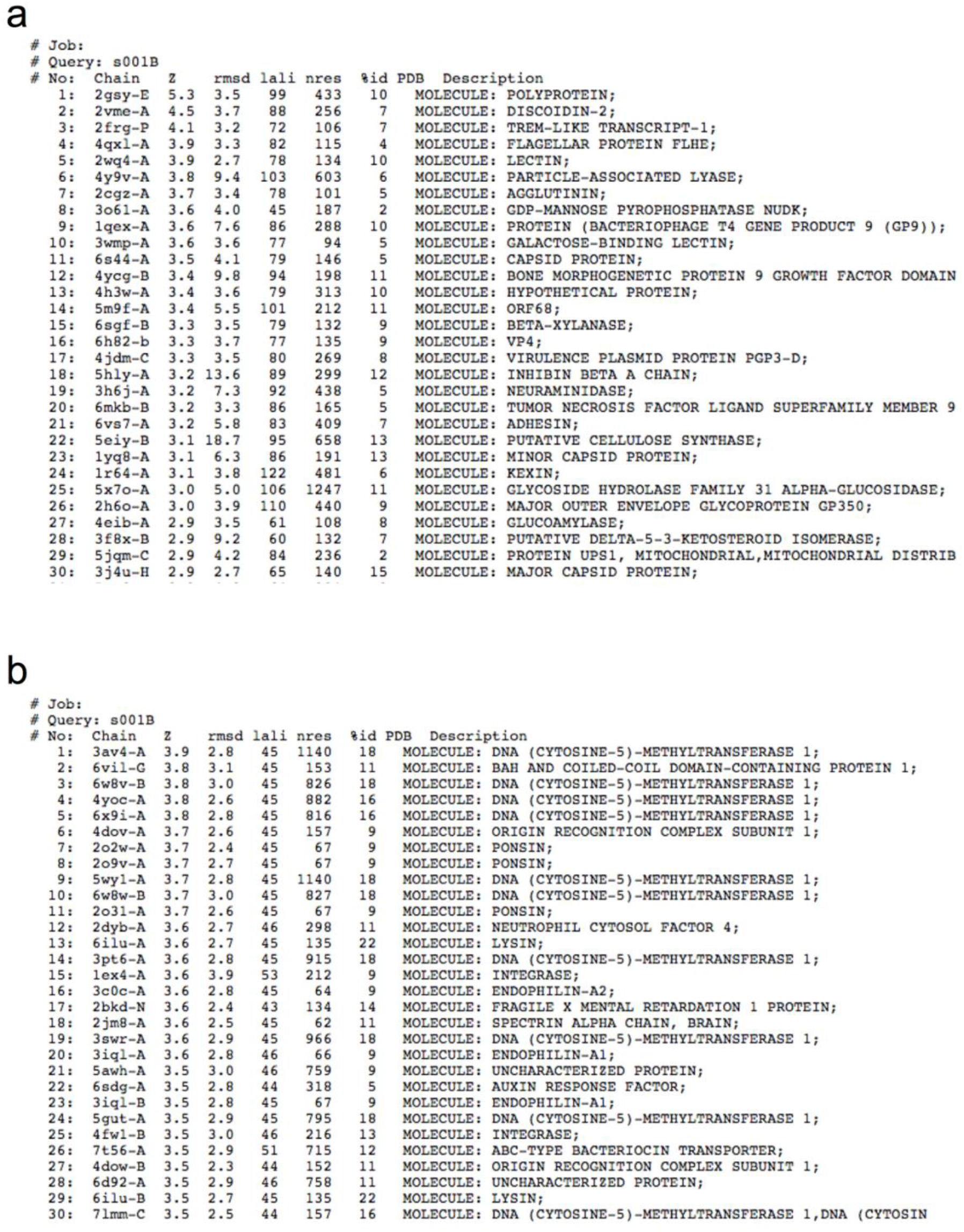
DALI analyses show low structural similarity of both pb5 and Llp to other proteins. Top 30 hits are shown for pb5 (**a**) and Llp (**b**). Coordinates were submitted to the webserver at http://ekhidna2.biocenter.helsinki.fi/dali/.

**Extended Data Figure 4.**
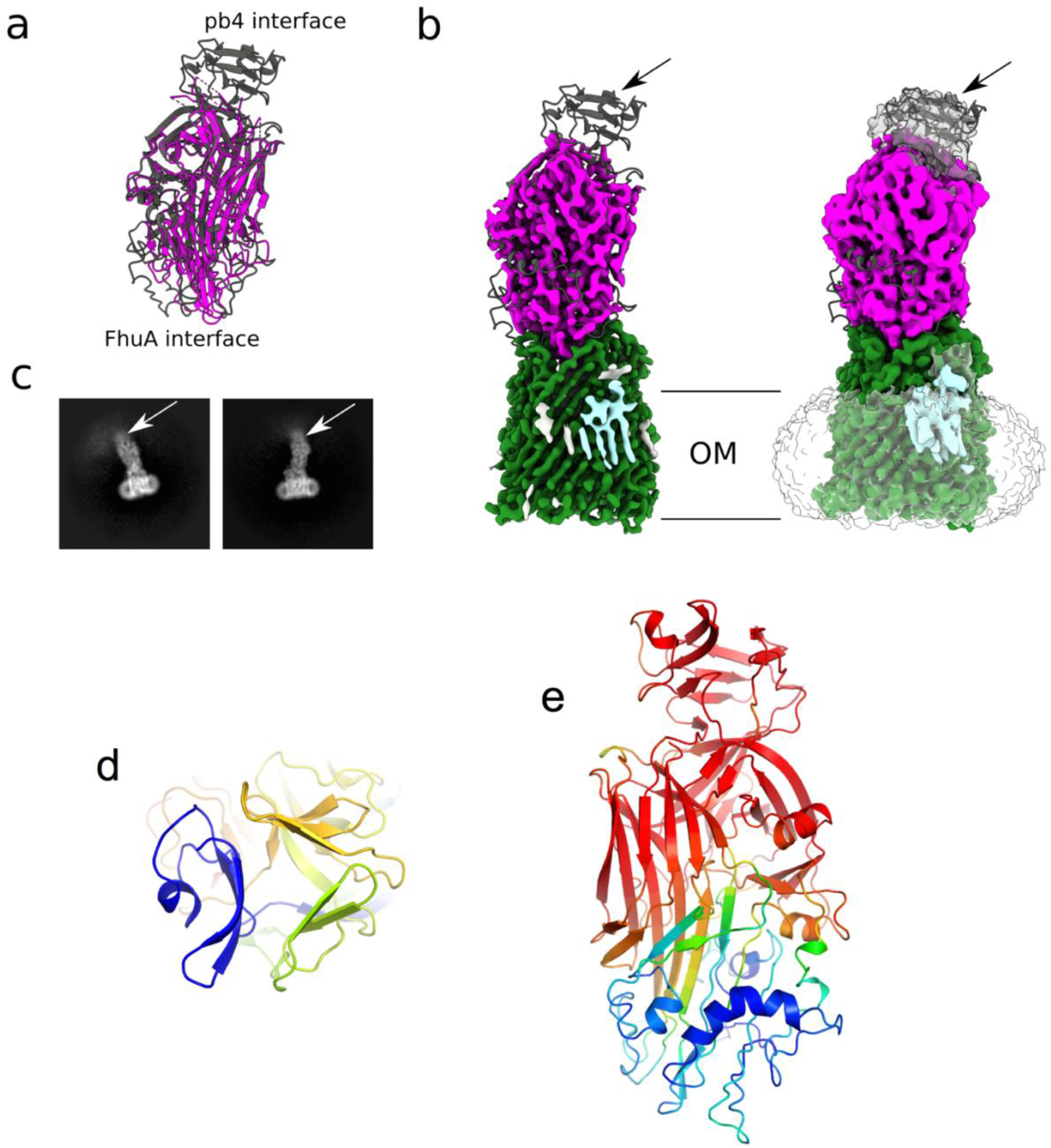
The pb4-interacting domain of pb5 is disordered in the FhuA-pb5 cryo-EM structure. **a**, Overlay of the AlphaFold2 prediction of pb5 (dark gray) with the experimental structure (magenta). **b**, Low threshold cryo-EM map on the right showing the pb4-interacting domain (arrow). Micelle density is transparent. **c**, Representative 2D classes for FhuA-pb5, showing fuzzy density for the pb4-interacting domain of pb5 (arrow). **d**, Top-down view of the pb4-interacting domain from the AlphaFold2 prediction, showing the pseudo 3-fold symmetry. e, Pb5 AlphaFold2 prediction coloured based on pLDDT confidence values (red, high; blue, low; minimum and maximum values are 24 and 97 respectively).

**Extended Data Figure 5.**
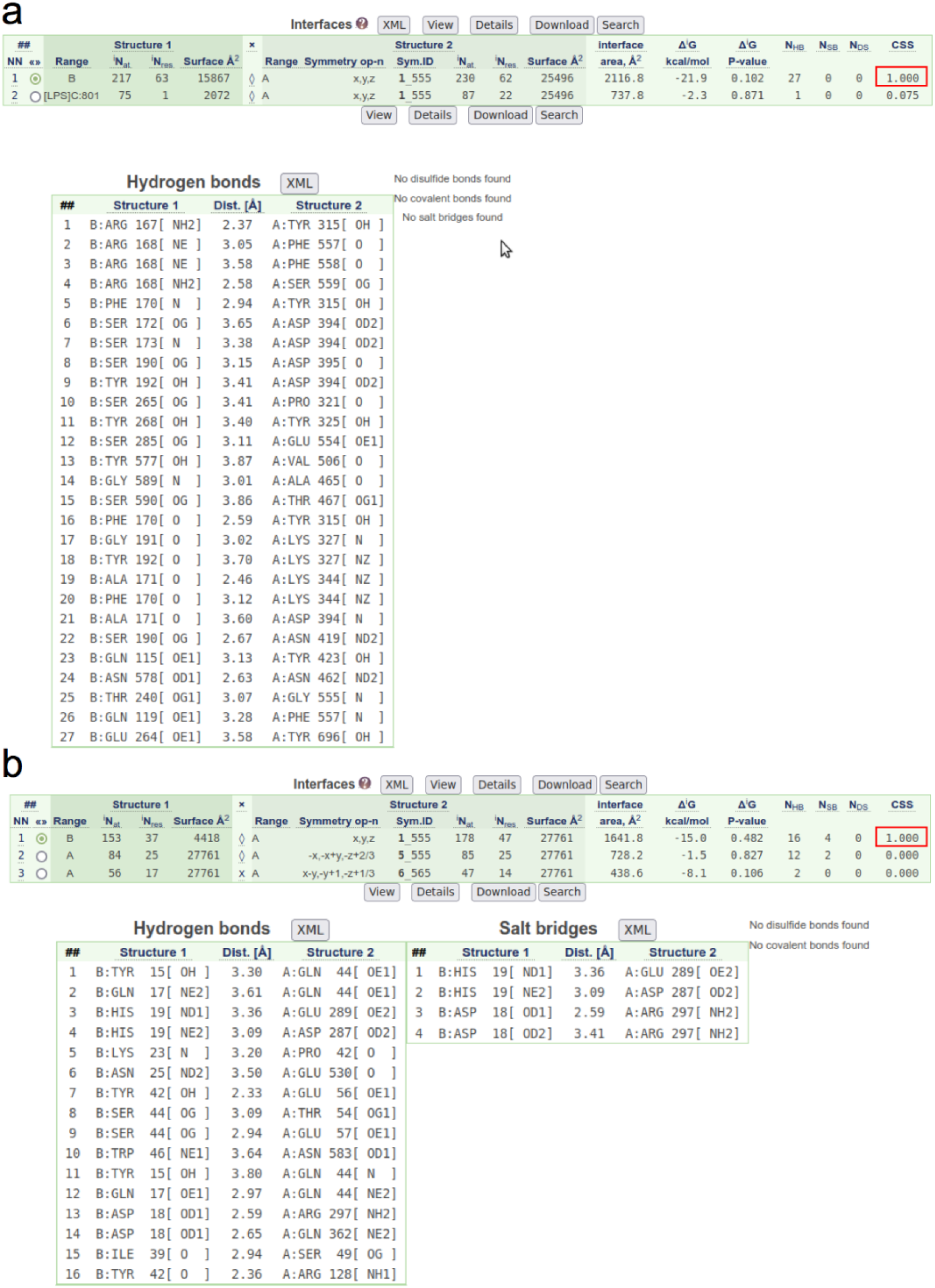
PISA analyses for FhuA-pb5 (**a**) and FhuA-Llp (**b**). The upper panels show an overview of the relevant interfaces. CSS stands for complexation significance score (maximum value 1.000). In both cases, FhuA is molecule A.

**Extended Data Figure 6.**
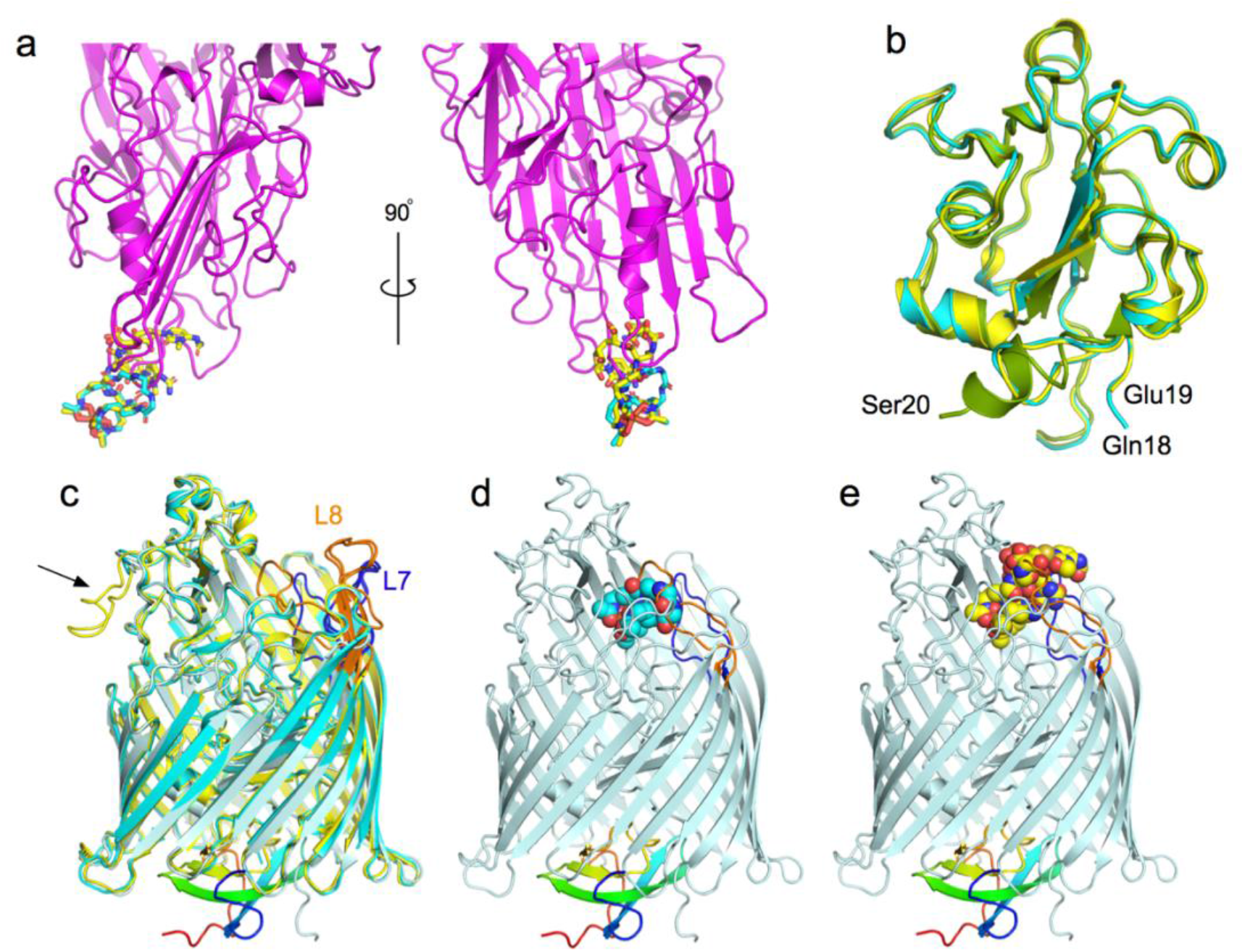
Pb5 and Llp binding to FhuA abolishes binding (and transport) of the ligands ferrichrome and albomycin. **a**, Cartoon model for FhuA-bound pb5 in two orientations, showing the overlap of pb5 loops with ferrichrome (cyan sticks) and albomycin (yellow sticks) bound to FhuA (PDB IDs 1BY5 and 1QKX respectively). **b**, Superpositions of the FhuA plug domains for FhuA-pb5 (green), FhuA-ferrichrome (cyan), and FhuA-albomycin (yellow). The N-terminal residues visible in the structures are labeled. **c**, Superposition of FhuA-Llp (light blue, with Llp in rainbow) with FhuA-ferrichrome (cyan) and FhuA-albomycin (yellow). For clarity, the ligands are not shown. Loops EL7 are EL8 are coloured blue and orange, respectively. **d**,**e** FhuA-Llp with superposed ferrichrome (**d**) or albomycin (**e**) shown as space-filling models. In both cases, the inward folded FhuA loops EL7 and EL8 clash with the ligands.

**Extended Data Figure 7.**
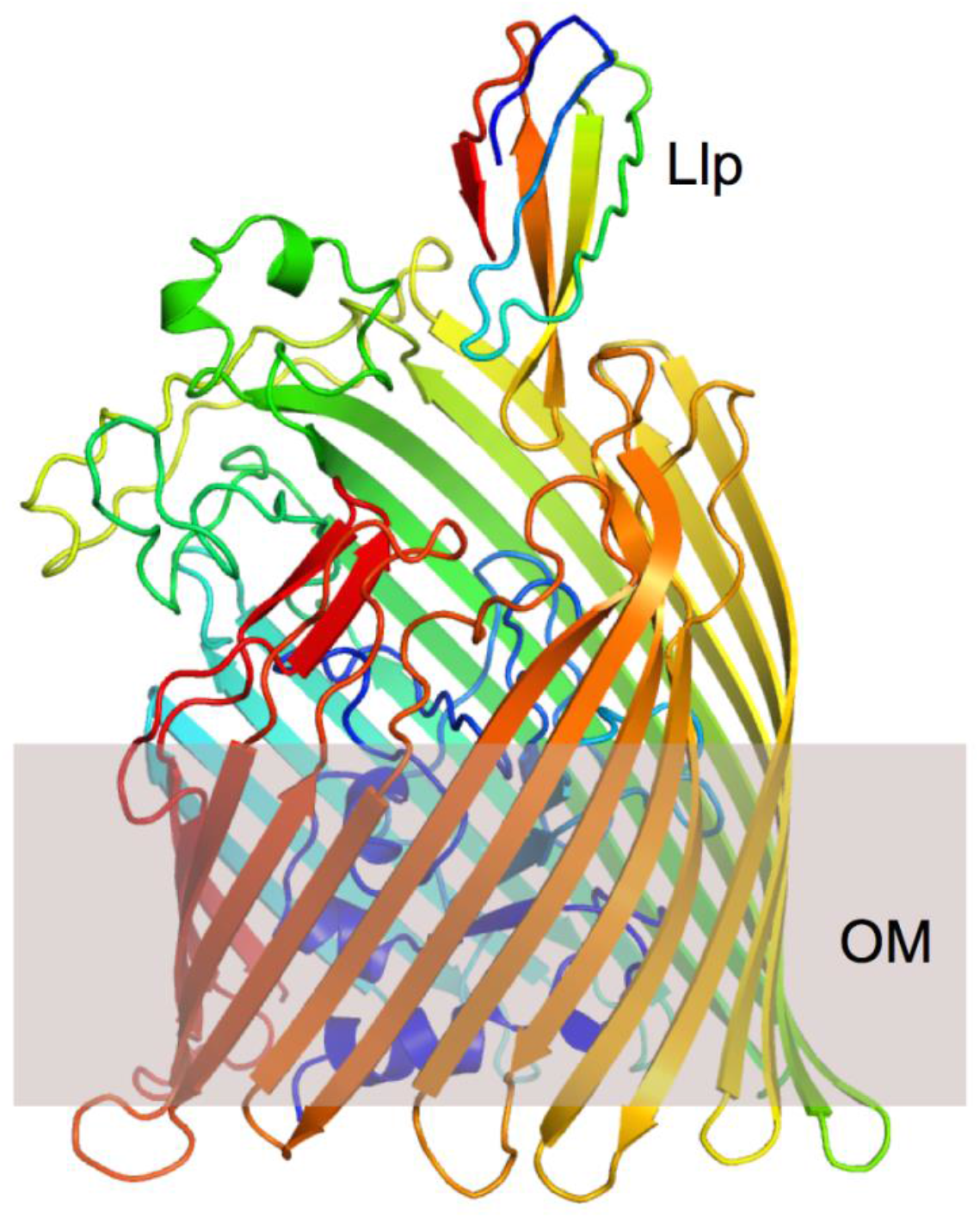
AlphaFold2 prediction of the FhuA-Llp complex. Both proteins are shown in rainbow representation (N-terminus, blue). The location of the hydrophobic core of the OM is indicated.

**Extended Data Figure 8.**
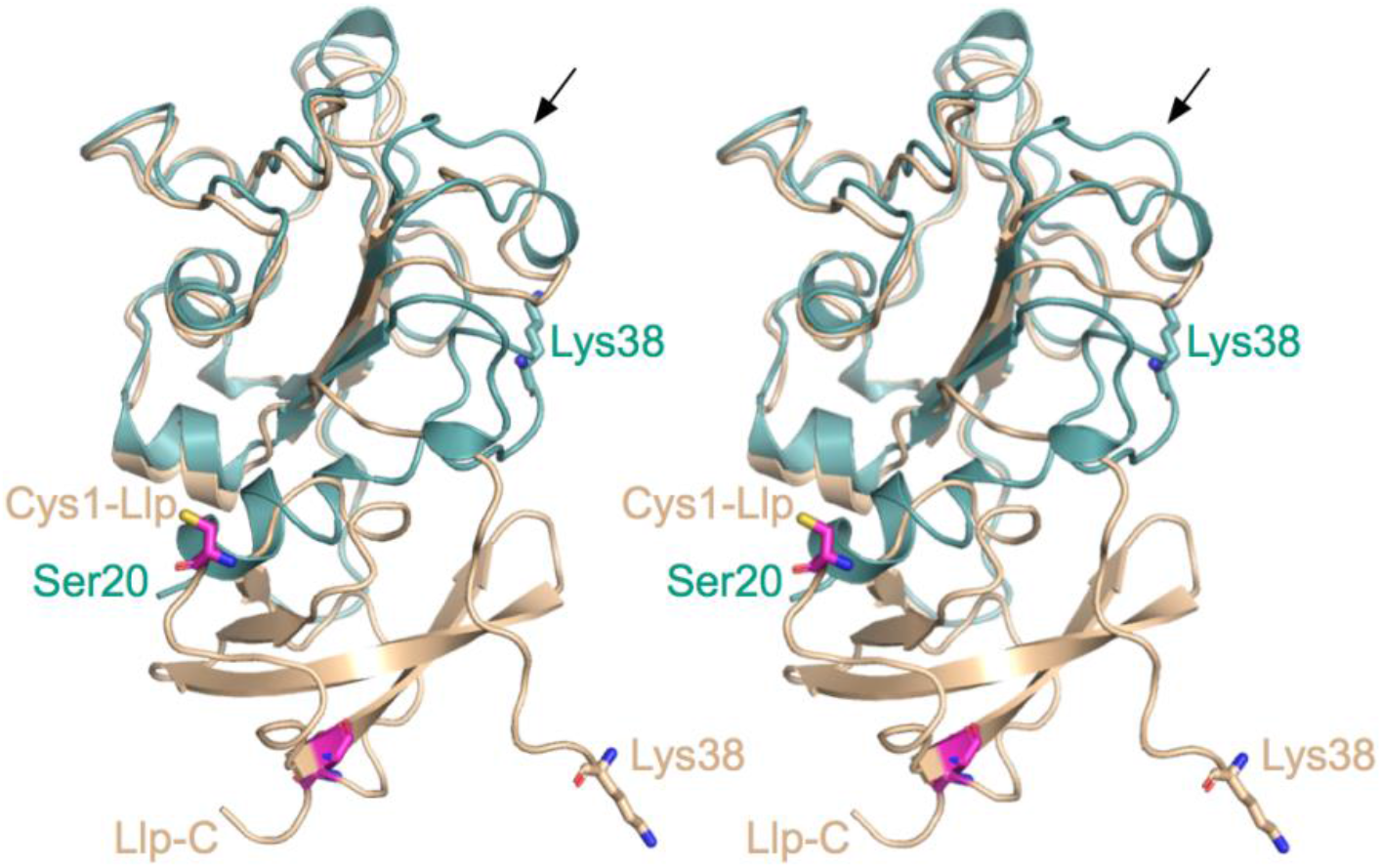
Conformational differences in the plug of FhuA-Llp. Stereo superposition of the free FhuA plug (teal) and the plug of FhuA-Llp (wheat). Cysteine residues of Llp are shown as magenta stick models. The visible N-termini are labeled and correspond to Ser20 in free FhuA and Lys38 in FhuA-Llp. The periplasmic space is at the bottom of the figure.

**Extended Data Figure 9.**
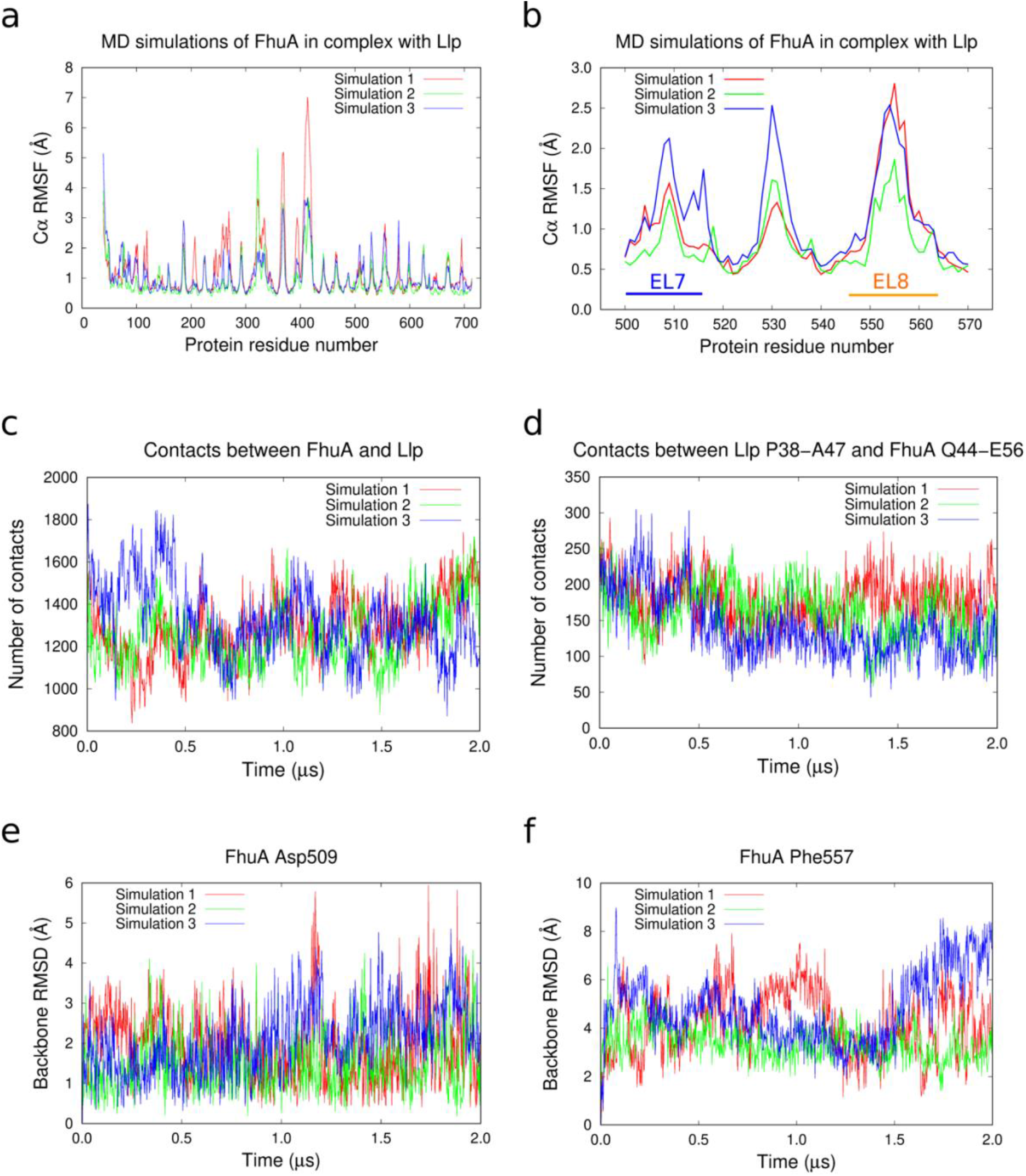
The FhuA-Llp interaction is stable and the conformations of EL7 and EL8 are not caused by crystal contacts. MD simulations showing Cα root-mean-square-fluctuations for FhuA (**a**) and the EL7-EL8 region (**b**). **c**,**d** Total number of contacts between FhuA and Llp (**c**) and between Llp Pro38-Ala47 and FhuA plug residues Gln44-Glu56 (**d**). A contact is defined as an atom-center to atom-center distance smaller than 4 Å, evaluated every 2 ns. **e**,**f** The positions of the tips of EL7 (Asp509; **e**) and EL8 (Phe557; **f**) are relatively stable during the 2 μs simulations compared to the starting structure.

**Extended Data Figure 10.**
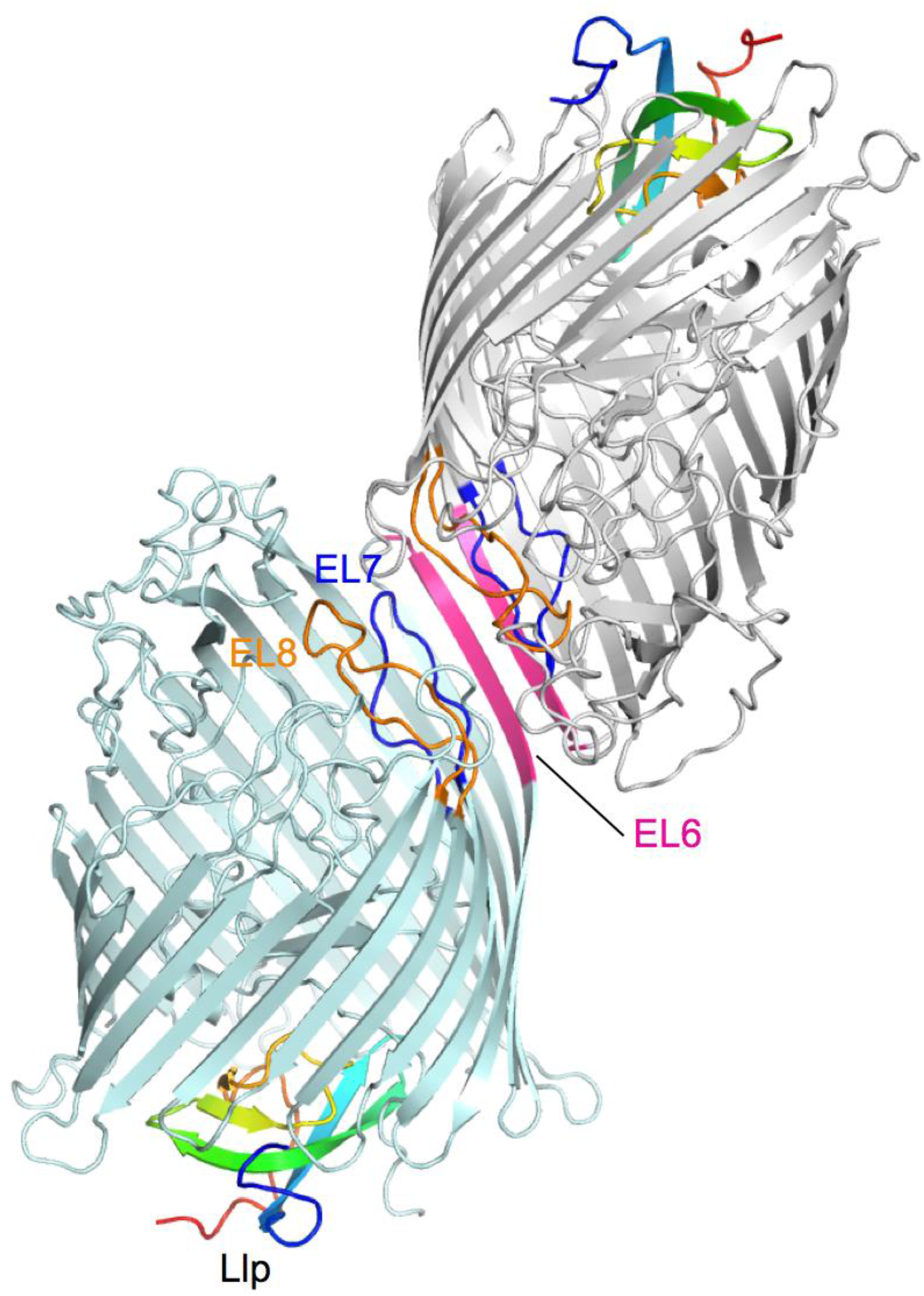
Crystal contacts in FhuA-Llp. Due to the inward-folded positions of EL7 and EL8, the exposed edge of one of the β-strands of EL6 is available to interact with a symmetry-related molecule (gray) via β-augmentation. This mode of crystal packing is not possible in free FhuA. MD simulations suggest that the conformational changes result in the observed crystal packing, and not the other way around.

**Supplementary Table 1.**
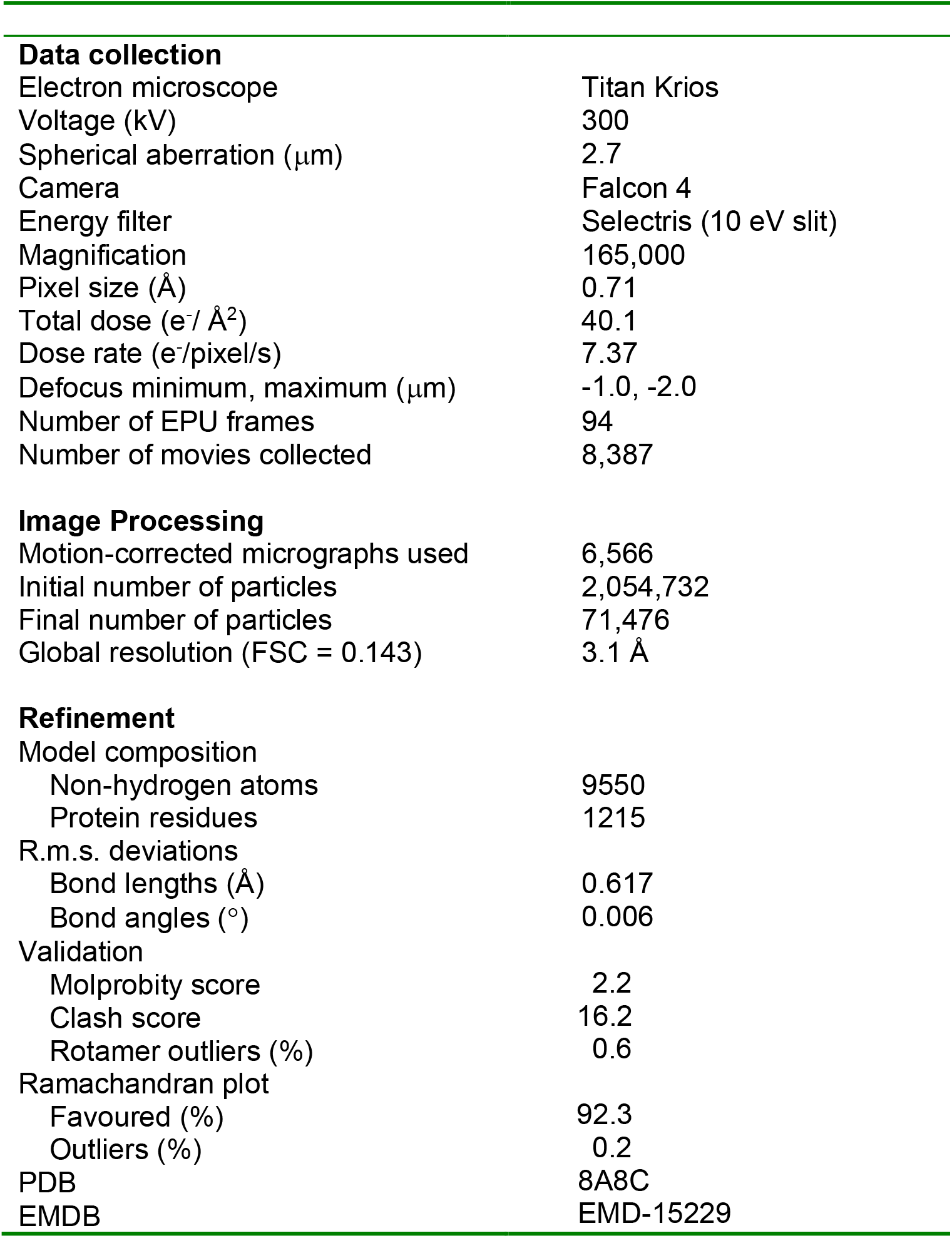
Cryo-EM data acquisition, processing and FhuA-pb5 model parameters.

**Supplementary Table 2.**
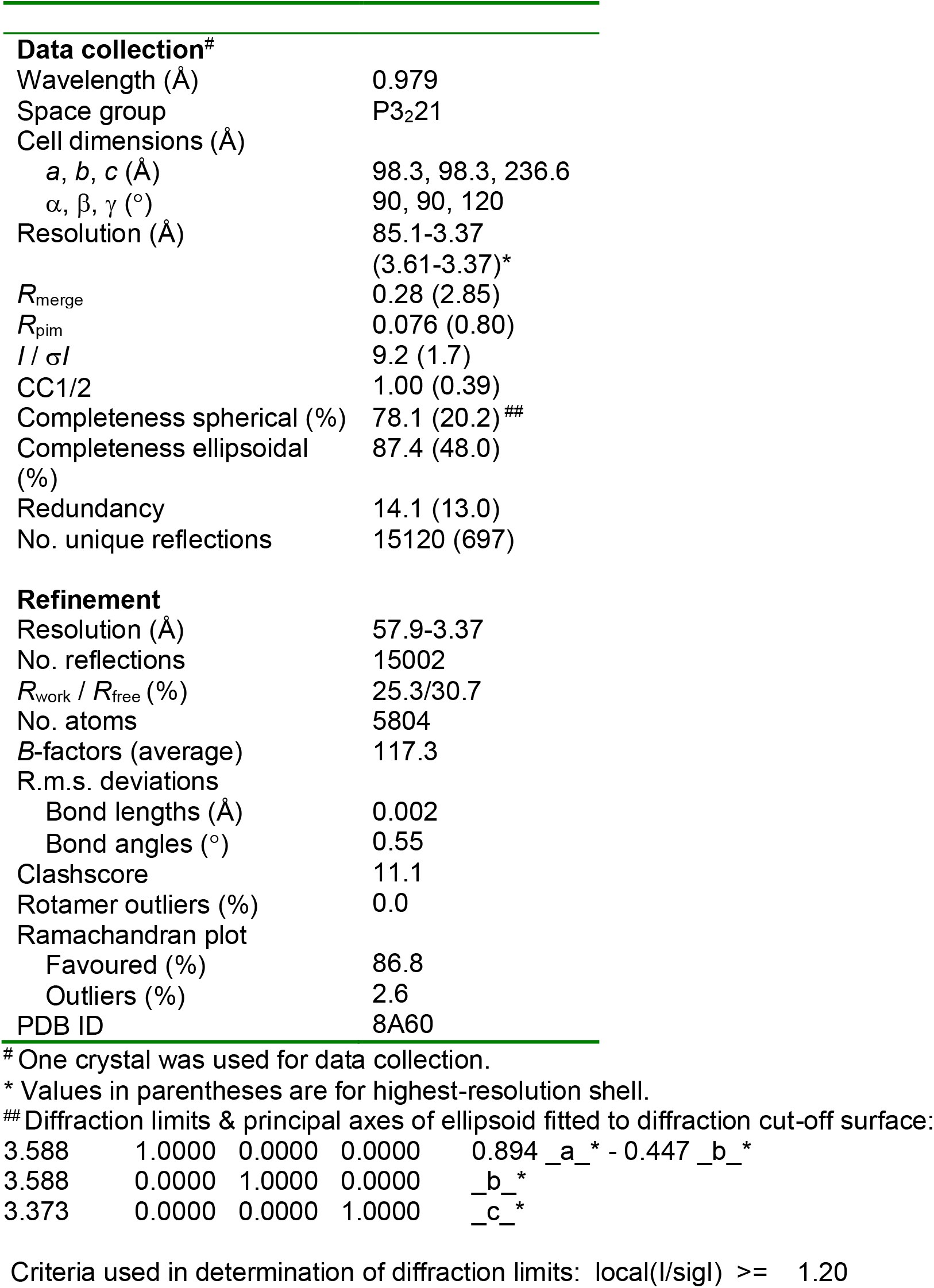
X-ray crystallographic data collection and refinement statistics for FhuA-Llp

|  |  |
| --- | --- |
| **Data collection<sup>#</sup>** |  |
| Wavelength (Å) | 0.979 |
| Space group | P3<sub>2</sub>21 |
| Cell dimensions (Å) |  |
| *a*, *b*, *c* (Å) | 98.3, 98.3, 236.6 |
| $\alpha$, $\beta$, $\gamma$ (°) | 90, 90, 120 |
| Resolution (Å) | 85.1-3.37 |
|  | (3.61-3.37)* |
| *R*<sub>merge</sub> | 0.28 (2.85) |
| *R*<sub>pim</sub> | 0.076 (0.80) |
| *I* / $\sigma$ *I* | 9.2 (1.7) |
| CC1/2 | 1.00 (0.39) |
| Completeness spherical (%) | 78.1 (20.2) ## |
| Completeness ellipsoidal (%) | 87.4 (48.0) |
| Redundancy | 14.1 (13.0) |
| No. unique reflections | 15120 (697) |
| **Refinement** |  |
| Resolution (Å) | 57.9-3.37 |
| No. reflections | 15002 |
| *R*<sub>work</sub> / *R*<sub>free</sub> (%) | 25.3/30.7 |
| No. atoms | 5804 |
| *B*-factors (average) | 117.3 |
| R.m.s. deviations |  |
| Bond lengths (Å) | 0.002 |
| Bond angles (°) | 0.55 |
| Clashscore | 11.1 |
| Rotamer outliers (%) | 0.0 |
| Ramachandran plot |  |
| Favoured (%) | 86.8 |
| Outliers (%) | 2.6 |
| PDB ID | 8A60 |

<sup>#</sup> One crystal was used for data collection.
\* Values in parentheses are for highest-resolution shell.

Diffraction limits & principal axes of ellipsoid fitted to diffraction cut-off surface:
|  |  |  |  |  |
| --- | --- | --- | --- | --- |
| 3.588 | 1.0000 | 0.0000 | 0.0000 | 0.894 _a_ * - 0.447 _b_ * |
| 3.588 | 0.0000 | 1.0000 | 0.0000 | _b_ * |
| 3.373 | 0.0000 | 0.0000 | 1.0000 | _c_ * |

Criteria used in determination of diffraction limits: local(*I*/ $\sigma$ *I*) ≥ 1.20

## Notes

### Competing Interest Statement

The authors have declared no competing interest.

## References

1. Nagel T, Musila L, Muthoni M, Nikolich M, Nakavuma JL, Clokie MR. Phage banks as potential tools to rapidly and cost-effectively manage antimicrobial resistance in the developing world. Curr Opin Virol. 53, 101208 (2022).

2. Meile S, Du J, Dunne M, Kilcher S, Loessner MJ. Engineering therapeutic phages for enhanced antibacterial efficacy. Curr Opin Virol. 52, 182–191 (2022).

3. Gurney J, Brown SP, Kaltz O, Hochberg ME. Steering Phages to Combat Bacterial Pathogens. Trends Microbiol. 28, 85–94 (2020).

4. Nobrega FL, Vlot M, de Jonge PA, Dreesens LL, Beaumont HJE, Lavigne R, Dutilh BE, Brouns SJJ. Targeting mechanisms of tailed bacteriophages. Nat Rev Microbiol. 16, 760–773 (2018).

5. Hutchison CA 3rd, Sinsheimer RL. Requirement of protein synthesis for bacteriophage phi X174 superinfection exclusion. J Virol. 8, 121–4 (1971).

6. Luria, S. E., and M. Delbruck.. Mutations of bacteria from virus sensitivity to virus resistance. Genetics 28, 491–511 (1943).

7. Keen EC. A century of phage research: bacteriophages and the shaping of modern biology. BioEssays 37, 6–9 (2015).

8. Demerec M, Fano U. Bacteriophage-Resistant Mutants in Escherichia Coli. Genetics 30, 119–36 (1945).

9. Wang J, Yan J, Vincent M, Sun Y, Yu H, Wang J, Bao Q, Kong H, Hu S. Complete genome sequence of bacteriophage T5. Virology 332, 45–65 (2005).

10. Effantin G, Boulanger P, Neumann E, Letellier L, Conway JF. Bacteriophage T5 structure reveals similarities with HK97 and T4 suggesting evolutionary relationships. J Mol Biol. 361, 993–1002 (2006).

11. Arnaud CA, Effantin G, Vivès C, Engilberge S, Bacia M, Boulanger P, Girard E, Schoehn G, Breyton C. Bacteriophage T5 tail tube structure suggests a trigger mechanism for Siphoviridae DNA ejection. Nat Commun. 8, 1953 (2017).

12. Boulanger P, Jacquot P, Plançon L, Chami M, Engel A, Parquet C, Herbeuval C, Letellier L. Phage T5 straight tail fiber is a multifunctional protein acting as a tape measure and carrying fusogenic and muralytic activities. J Biol Chem. 283, 13556–64 (2008).

13. Zivanovic Y, Confalonieri F, Ponchon L, Lurz R, Chami M, Flayhan A, Renouard M, Huet A, Decottignies P, Davidson AR, Breyton C, Boulanger P. Insights into bacteriophage T5 structure from analysis of its morphogenesis genes and protein components. J Virol. 88, 1162–74 (2014).

14. Heller K, Braun V. Accelerated adsorption of bacteriophage T5 to Escherichia coli F, resulting from reversible tail fiber-lipopolysaccharide binding. J Bacteriol. 139, 32–8 (1979).

15. Heller KJ. Identification of the phage gene for host receptor specificity by analyzing hybrid phages of T5 and BF23. Virology 139, 11–21 (1984).

16. Braun, V., K. Schaller, and H. Wolf. A common receptor protein for phage T5 and colicin M in the outer membrane of Escherichia coli B. Biochim. Biophys. Acta 323, 87–97 (1973).

17. Braun V. FhuA (TonA), the career of a protein. J Bacteriol. 191, 3431–6 (2009).

18. Flayhan A, Wien F, Paternostre M, Boulanger P, Breyton C. New insights into pb5, the receptor binding protein of bacteriophage T5, and its interaction with its Escherichia coli receptor FhuA. Biochimie. 94, 1982–9 (2012).

19. Plançon L, Janmot C, le Maire M, Desmadril M, Bonhivers M, Letellier L, Boulanger P. J Characterization of a high-affinity complex between the bacterial outer membrane protein FhuA and the phage T5 protein pb5. Mol Biol. 318, 557–69 (2002).

20. Breyton C, Flayhan A, Gabel F, Lethier M, Durand G, Boulanger P, Chami M, Ebel C. Assessing the conformational changes of pb5, the receptor-binding protein of phage T5, upon binding to its Escherichia coli receptor FhuA. J Biol Chem. 288, 30763–30772 (2013).

21. Veesler D, Cambillau C. A common evolutionary origin for tailed-bacteriophage functional modules and bacterial machineries. Microbiol Mol Biol Rev. 75, 423–33 (2011).

22. Decker K, Krauel V, Meesmann A, Heller KJ. Lytic conversion of Escherichia coli by bacteriophage T5: blocking of the FhuA receptor protein by a lipoprotein expressed early during infection. Mol Microbiol. 12, 321–32 (1994).

23. Braun V, Killmann H, Herrmann C. Inactivation of FhuA at the cell surface of Escherichia coli K-12 by a phage T5 lipoprotein at the periplasmic face of the outer membrane. J Bacteriol. 176, 4710–7 (1994).

24. Pedruzzi I, Rosenbusch JP, Locher KP. Inactivation in vitro of the Escherichia coli outer membrane protein FhuA by a phage T5-encoded lipoprotein. FEMS Microbiol Lett. 168, 119–25 (1998).

25. Wietzorrek A, Schwarz H, Herrmann C, Braun V. The genome of the novel phage Rtp, with a rosette-like tail tip, is homologous to the genome of phage T1. J Bacteriol. 188, 1419–36 (2006).

26. Locher KP, Rees B, Koebnik R, Mitschler A, Moulinier L, Rosenbusch JP, Moras D. Transmembrane signaling across the ligand-gated FhuA receptor: crystal structures of free and ferrichrome-bound states reveal allosteric changes. Cell 95, 771–8 (1998).

27. Jumper J, et al. Highly accurate protein structure prediction with AlphaFold. Nature 596, 583–589 (2021).

28. Varadi M, et al. AlphaFold Protein Structure Database: massively expanding the structural coverage of protein-sequence space with high-accuracy models. Nucleic Acids Res. 50, D439–D444 (2022).

29. Ferguson AD, Hofmann E, Coulton JW, Diederichs K, Welte W.Siderophore-mediated iron transport: crystal structure of FhuA with bound lipopolysaccharide. Science 282, 2215–20 (1998).

30. Holm L. Using Dali for protein structure comparison. Methods Mol. Biol. 2112, 29–42 (2020).

31. Mondigler M, Holz T, Heller KJ. Identification of the receptor-binding regions of pb5 proteins of bacteriophages T5 and BF23. Virology 219, 19–28 (1996).

32. Krissinel E, Henrick K. Inference of macromolecular assemblies from crystalline state. J Mol Biol. 372, 774–97 (2007).

33. Endriss F, Braun V. Loop deletions indicate regions important for FhuA transport and receptor functions in Escherichia coli. J Bacteriol. 186, 4818–23 (2004).

34. Ferguson AD, Braun V, Fiedler HP, Coulton JW, Diederichs K, Welte W. Crystal structure of the antibiotic albomycin in complex with the outer membrane transporter FhuA. Protein Sci. 9, 956–63 (2000).

35. Pawelek PD, Croteau N, Ng-Thow-Hing C, Khursigara CM, Moiseeva N, Allaire M, Coulton JW. Structure of TonB in complex with FhuA, E. coli outer membrane receptor. Science 312, 1399–402 (2006).

36. Maffei E, Shaidullina A, Burkolter M, Heyer Y, Estermann F, Druelle V, Sauer P, Willi L, Michaelis S, Hilbi H, Thaler DS, Harms A. Systematic exploration of Escherichia coli phage-host interactions with the BASEL phage collection. PLoS Biol. 19, e3001424 (2021).

37. Shi K, Oakland JT, Kurniawan F, Moeller NH, Banerjee S, Aihara H.Structural basis of superinfection exclusion by bacteriophage T4 Spackle. Commun Biol. 3, 691 (2020).

38. Cowles CE, Li Y, Semmelhack MF, Cristea IM, Silhavy TJ. The free and bound forms of Lpp occupy distinct subcellular locations in Escherichia coli. Mol Microbiol. 79, 1168–81 (2011).

39. Bonhivers M, Ghazi A, Boulanger P, Letellier L. FhuA, a transporter of the Escherichia coli outer membrane, is converted into a channel upon binding of bacteriophage T5. EMBO J. 15, 1850–6 (1996).

40. Bonhivers M, Plançon L, Ghazi A, Boulanger P, le Maire M, Lambert O, Rigaud JL, Letellier L.FhuA, an Escherichia coli outer membrane protein with a dual function of transporter and channel which mediates the transport of phage DNA. Biochimie. 80, 363–9 (1998).

41. Böhm J, Lambert O, Frangakis AS, Letellier L, Baumeister W, Rigaud JL.FhuA-mediated phage genome transfer into liposomes: a cryo-electron tomography study. Curr Biol. 11, 1168–75 (2002).

42. Zampara A, Sørensen MCH, Grimon D, Antenucci F, Vitt AR, Bortolaia V, Briers Y, Brøndsted L. Exploiting phage receptor binding proteins to enable endolysins to kill Gram-negative bacteria. Sci Rep. 10, 12087 (2020).

43. Punjani, A., Rubinstein, J.L., Fleet, D.J. & Brubaker, M.A. cryoSPARC: algorithms for rapid unsupervised cryo-EM structure determination. Nature Methods 14, 290–296 (2017).

44. Punjani, A., Zhang, H. & Fleet, D.J. Non-uniform refinement: adaptive regularization improves single-particle cryo-EM reconstruction. Nat Methods 17, 1214–1221 (2020).

45. Adams PD et al. PHENIX: a comprehensive Python-based system for macromolecular structure solution. Acta Crystallogr D Biol Crystallogr 66, 213–21 (2010).

46. Emsley P & Cowtan K. Coot: model-building tools for molecular graphics. Acta Crystallogr D Biol Crystallogr 60, 2126–32 (2004).

47. McCoy AJ et al. Phaser crystallographic software. J Appl Crystallogr 40, 658–674 (2007).

48. Vonrhein C, Flensburg C, Keller P, Sharff A, Smart O, Paciorek W, Womack T, Bricogne G. Data processing and analysis with the autoPROC toolbox. Acta Cryst. D67, 293–302 (2011).

49. Tickle IJ, Flensburg C, Keller P, Paciorek W, Sharff A, Vonrhein C, Bricogne G. STARANISO. Cambridge, United Kingdom: Global Phasing Ltd (2018).

50. S. Jo, T. Kim, V.G. Iyer, and W. Im. CHARMM-GUI: A Web-based Graphical User Interface for CHARMM. J. Comput. Chem. 29, 1859–1865 (2008). DOI: 10.1002/jcc.20945.

51. H. J. C. Berendsen, J. P. M. Postma, W. F. van Gunsteren, A. DiNola, and J. R. Haak. Molecular dynamics with coupling to an external bath. J. Chem. Phys. 81, 3684–3690 (1984).

52. S. Nosé. A molecular dynamics method for simulations in the canonical ensemble. Mol. Phys. 52, 255–268 (1984).

53. W. G. Hoover. Canonical Dynamics: Equilibrium Phase-Space Distributions. Phys. Rev. A, 31, 1695–1697 (1985).

54. M. Parrinello and A. Rahman. Polymorphic transitions in single crystals: A new molecular dynamics method. J. Appl. Phys. 52, 7182–7190 (1981).

55. B. Hess, H. Bekker, H. J. C. Berendsen, and J. G. E. M. Fraaije. LINCS: A Linear Constraint Solver for Molecular Simulations. J. Comput. Chem. 18, 1463–1472 (1998).

56. U. Essmann, L. Perera, M. L. Berkowitz, T. Darden, H. Lee, and L. G. Pedersen. A smooth particle mesh Ewald method. J. Chem. Phys. 103, 8577–8593 (1995).

57. M. J. Abraham et al. GROMACS: High performance molecular simulations through multi-level parallelism from laptops to supercomputers. SoftwareX 1-2, 19–25 (2015).

58. J. Huang et al. CHARMM36m: An Improved Force Field for Folded and Intrinsically Disordered Proteins. Nat. Methods 14, 71–73 (2016).

59. J. B. Klauda et al. Update of the CHARMM All-Atom Additive Force Field for Lipids: Validation on Six Lipid Types. J. Phys. Chem. B 114, 7830–7843 (2010).

60. W. Humphrey, A. Dalke, and K. Schulten. VMD - Visual Molecular Dynamics. J. Molec. Graphics 14, 33–38 (1996).

